# Multi-Modal Profiling of Human Fetal Liver-Derived Hematopoietic Stem Cells Reveals the Molecular Signature of Engraftment Potential

**DOI:** 10.1101/2020.11.11.378620

**Authors:** Kim Vanuytsel, Carlos Villacorta-Martin, Jonathan Lindstrom-Vautrin, Zhe Wang, Wilfredo F. Garcia-Beltran, Vladimir Vrbanac, Taylor M. Matte, Todd W. Dowrey, Sara S. Kumar, Mengze Li, Ruben Dries, Joshua D. Campbell, Anna C. Belkina, Alejandro B. Balazs, George J. Murphy

## Abstract

The human hematopoietic stem cell (HSC) harbors remarkable regenerative potential that can be harnessed therapeutically. During early development, HSCs in the fetal liver (FL) undergo active expansion while simultaneously retaining robust engraftment capacity, yet the underlying molecular program responsible for their efficient engraftment remains unclear. We profiled 26,407 FL cells at both transcriptional and protein levels including over 7,000 highly enriched and functional FL HSCs to establish a detailed molecular signature of engraftment potential. Integration of transcript and linked cell surface marker expression revealed a generalizable signature defining functional FL HSCs and allowed for the stratification of enrichment strategies with high translational potential. This comprehensive, multi-modal profiling of engraftment capacity connects a critical biological function at a key developmental timepoint with its underlying molecular drivers, serving as a useful resource for the field.

## INTRODUCTION

The human hematopoietic stem cell (HSC) has been the focus of intense study due to its remarkable regenerative potential and its therapeutic utility in treating a variety of diseases. HSCs reside at the apex of the hematopoietic system and have the capacity to both self-renew and differentiate into all mature blood cell types. As such, these cells, and the hierarchical structure of their progeny, represent an ideal system in which to ask fundamental biological questions and discover key insights into hematopoietic development and disease. These insights could be harnessed to improve *ex vivo* expansion methods, increase engraftment efficiency, and enable *in vitro* HSC generation from pluripotent stem cell (PSC) sources.

The first HSCs emerge in the aorta-gonad-mesonephros (AGM) region prior to travelling to the fetal liver (FL), where they undergo expansion to create the HSC pool that sustains hematopoiesis for the lifetime of an individual. The FL remains the main site of hematopoiesis until HSCs colonize their final niche in the bone marrow (BM) closer to birth (Ciriza et al., 2013; Holt and Jones, 2000) This dynamic transition is coupled to a switch from a proliferative to a predominantly quiescent phenotype postnatally (Ciriza et al., 2013), reflecting just one of several differences between developmentally distinct HSCs (McKinney-Freeman et al., 2012). A key functional difference between FL-derived and more mature HSCs is the superior engraftment potential of FL when compared to cord blood (CB) and BM cells (Holyoake et al., 1999). This retention of robust engraftment potential while undergoing active expansion highlights a unique feature of FL HSCs, given that proliferation and HSC functionality are inversely correlated postnatally (Rodriguez-Fraticelli et al., 2020; Walter et al., 2015). This prompted us to specifically profile FL-derived HSCs to establish a detailed molecular signature of engraftment potential.

To dissect the molecular underpinnings of engraftment potential at the highest possible resolution, we chose to combine several orthogonal single cell profiling approaches. The emergence of single cell RNA sequencing (scRNAseq) technologies has led to significant advances in terms of characterization of the larger hematopoietic stem and progenitor cell (HSPC) pool. Transcriptomic profiling of large numbers of single CD34^+^ cells has resulted in a more nuanced understanding of the postnatal HSPC compartment and has revealed a continuum of transcriptional states, rather than a succession of clearly demarcated progenitor stages (Belluschi et al., 2018; Pellin et al., 2019; Velten et al., 2017; Weinreb et al., 2020). Recent work in the context of the developmental cell atlas has extended such large-scale single cell transcriptomic profiling efforts to the prenatal stage, analyzing the dynamic hematopoietic composition of the entire FL and other tissues throughout early human development (Popescu et al., 2019). However, despite interrogation of large numbers of total cells, their relative scarcity resulted in the profiling of few truly functional HSCs. In this study, we focus specifically on this rare HSC fraction by profiling a highly enriched population of FL HSCs, which we confirm to be functional for engraftment, at both a transcriptional as well as protein level.

Several strategies exist to enrich for functional HSCs within a pool of HSPCs (Notta et al., 2011; Prashad et al., 2015; Subramaniam et al., 2019; Sumide et al., 2018), with a combination of GPI-80 and CD133 resulting in one of the highest frequencies described (~1/5) (Sumide et al., 2018). Glycophosphatidylinositol-anchored surface protein GPI-80 has been shown to mark a subpopulation of FL HSCs that combines self-renewal ability and engraftment potential (Prashad et al., 2015). Using this marker as the basis for functional HSC enrichment, we single cell profiled these highly-enriched FL cells to uncover the detailed molecular signature of engraftable FL HSCs. Further dissection of this engraftment profile revealed signatures highlighting the importance of proteome integrity maintenance as well as the prominent expression of factors linked to aging and the concomitant decline of HSC functionality. This comprehensive characterization of the engraftment signature of FL HSCs using multi-modal profiling to define an essential biological function at a key developmental timepoint will serve as a useful resource for the field. Our interactive dataset is freely available to all of the scientific community through the following platform: https://engraftable-hsc.cells.ucsc.edu

## RESULTS

### The hematopoietic landscape of human fetal liver at single cell resolution

To obtain a detailed molecular signature of human FL HSCs, we performed CITE-seq (Stoeckius et al., 2017), a technique that allows for the simultaneous assessment of transcript and cell surface marker level expression by combining droplet based single cell RNA sequencing (scRNAseq) and oligo-tagged antibodies. Following dissociation of human FL, cells were divided into either CD34^+^-enriched cells or CD34^−^ flowthrough cells via magnetic bead separation (**Figure S1A**). The CD34^−^ live cell fraction was further subdivided into GYPA^+^ and GYPA^−^ via fluorescence activated cell sorting (FACS) to capture populations of maturing erythroid progenitors that constitute a sizeable portion of the FL at this stage in development (CD34^−^GYPA^+^). From the CD34^+^-enriched fraction live CD34^+^ cells were sorted (CD34^+^bulk) to reflect the cell population that is routinely used in a clinical HSC transplantation setting. In a separate sample, we further enriched this population using FACS based on GPI-80 expression (GPI-80^+^), a marker tightly linked to engraftment potential (Prashad et al., 2015; Sumide et al., 2018), to focus our analysis on HSCs capable of long-term engraftment. Prior to sorting, cells were stained with a panel of oligo-tagged antibodies (**Table S1**) so that the antibody-derived tags (ADTs) corresponding to the cell surface markers present on each cell would also be captured in the sequencing data. Following data processing and quality control, this resulted in a total of 26,407 FL cells profiled, divided across the following samples: 8735 CD34^+^bulk cells, 7235 GPI-80^+^ cells, 6793 CD34^−^GYPA^−^ cells and 3644 CD34^−^GYPA^−^ cells (**Figure S1A**).

Transcriptomic analysis of all four samples combined resulted in an overview of the hematopoietic landscape at this developmental stage (**Figure 1A**). **Figure 1B** displays the distribution of the four samples within the combined data set, illustrating that the CD34^+^ HSC/multipotent progenitor (MPP) fraction represents cells belonging to the CD34^+^ bulk and the GPI-80^+^-enriched fraction. Greater than half of all of the assayed cells represented CD34^+^ HSCs/MPPs (**Figure 1A-E**). This point is reflected by both the mRNA expression pattern of *CD34* (**Figure 1C)** as well as the overlay of a previously established HSC/MPP signature within the human FL (**Figure 1D**) (Popescu et al., 2019) onto our combined data set (**Figure 1E**). Taking these CD34^+^ HSCs/MPPs as a starting point, gene expression changes over pseudotime were assessed for each major hematopoietic lineage (**Figure S1B-E**). This analysis confirmed downregulation of HSC/MPP marker genes and upregulation of key lineage identity genes as commitment progresses, reflecting the expected hematopoietic cell types at this developmental stage. More specifically, cells along the erythroid lineage trajectory upregulated *LSMD1* and *GATA2* in the erythro-megakaryo-mast progenitor (EMMP) stage and went on to express globin genes, *ITGA2B*, and *LMO4* upon further erythroid, megakaryocyte and mast cell commitment respectively (**Figure 1C, Figure S2B**). Cells along the lymphoid B cell trajectory subsequently upregulated *IGLL1*, *VPREB1* and *VPREB3* before expressing *CD79A* and *CD79B* as fully committed B cells (**Figure 1C, Figure S2C**). Cells specifying towards T and NK cell fates exhibited upregulation of *IL7R* and components of the T cell receptor CD3 complex (*CD3G*) in case of early lymphoid/ T cell commitment and expressed *NKG7*, *GZMA* and *TIGIT* upon NK cell specification (**Figure 1C, Figure S2D**). The myeloid trajectory was associated with upregulation of *MPO* early on as cells progressed through a neutrophil/myeloid progenitor stage. Further myeloid specification resulted in upregulation of *CST3*, *CSF1R*, *CD14* and *FCGR3A* (*CD16*) (**Figure 1C, Figure S2E**) and cells with signatures corresponding to macrophages, Kupffer cells and dendritic cells. As expected, cells specifying towards an erythroid fate downregulated *PTPRC* (*CD45*) expression, while myeloid and lymphoid specifying cells showed clear upregulation of this marker at the end of their differentiation trajectories (**Figure 1C, Figure S2B-E**). Interestingly, GYPA^+^ sorting also appeared to enrich for cells with a signature of B cells, consistent with earlier reports of *GYPA* expression in B cells (**Figure 1B,C, Figure S2C**) (Monaco et al., 2019).

**Figure 1.**
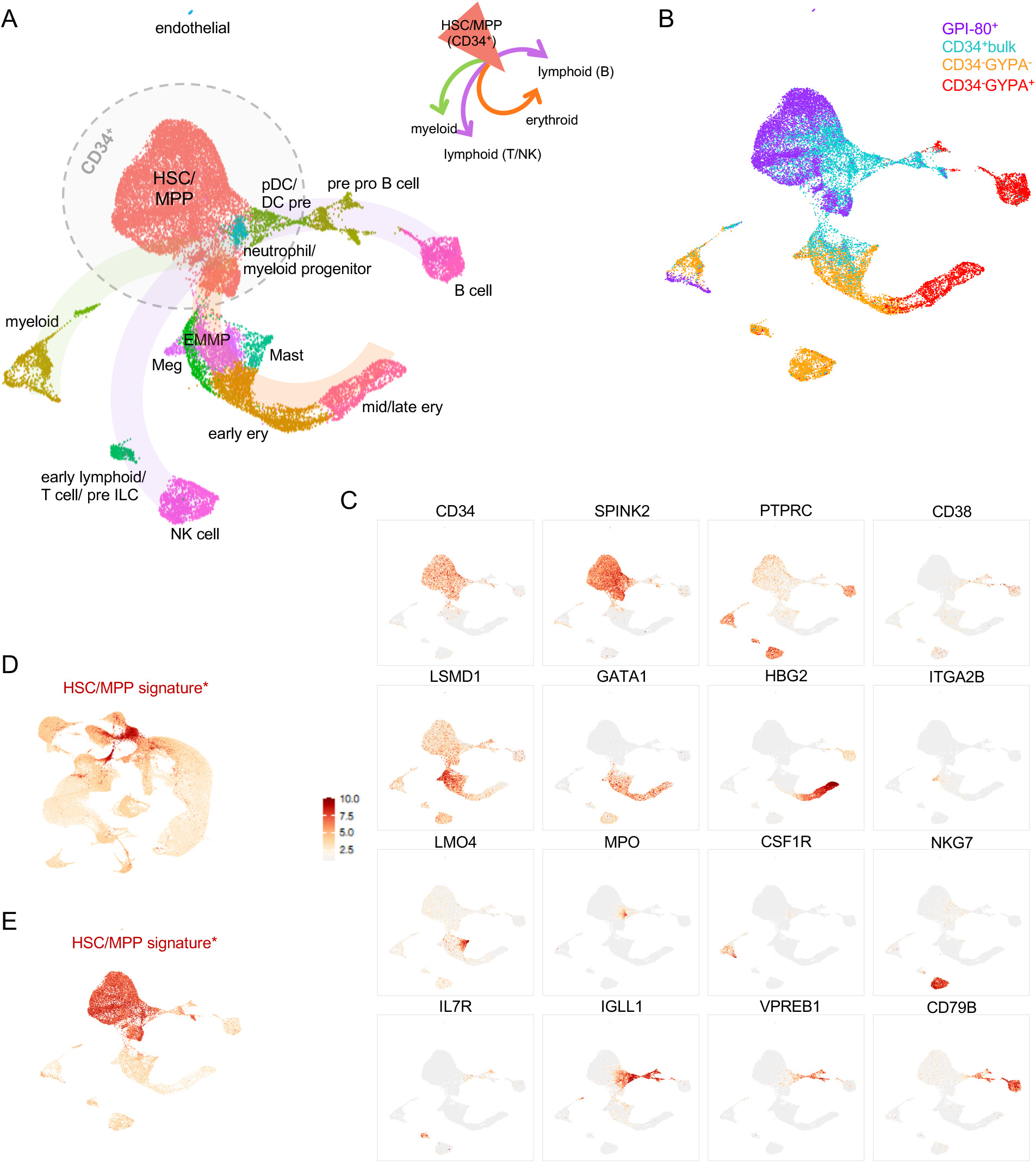
The hematopoietic landscape of human fetal liver at single cell resolution. (A) UMAP representation of the transcriptomic analysis of all 4 samples combined showing the cell types identified in each cluster. Cells are colored by cell lineage annotation, developed using the expression of key markers and DEG from the transcriptomic analysis. (B) UMAP representing the distribution of each sample within the combined analysis. Cells are colored by the sample origin of each cell. (C) UMAP overlays illustrating mRNA expression patterns of a set of HSC/MPP and lineage markers. The scale bar applies to the scaled expression shown in panels C-E. (D) Re-analysis of data from Popescu et al. (2019) to illustrate the identified HSC/MPP signature in its original context (Popescu et al., 2019). (E) Overlay of HSC/MPP signature identified in Popescu et al. (2019) onto the combined transcriptomic analysis described in this study.

### Single cell transcriptomic profiling of the engraftment potential of FL-derived HSCs

In parallel with capture on the 10X Genomics platform, cells from these same sorted fractions (with the exception of the CD34^−^GYPA^+^ fraction) were used in simultaneous transplantation experiments to assess engraftment capacity. These experiments revealed superior per-cell engraftment potential of the GPI-80^+^ fraction as compared to CD34^+^bulk and CD34^−^GYPA^−^ fractions, confirming enrichment for bona fide functional HSCs in this population (**Figure 2A**). Although the GPI-80^+^ fraction represented only 2.37% of the cells within the CD34^+^bulk population (**Figure S1A**), sorting and enrichment of this fraction enabled the analysis of 7235 GPI-80^+^ cells, resulting in profiling of the engraftable FL HSC at unprecedented resolution.

**Figure 2.**
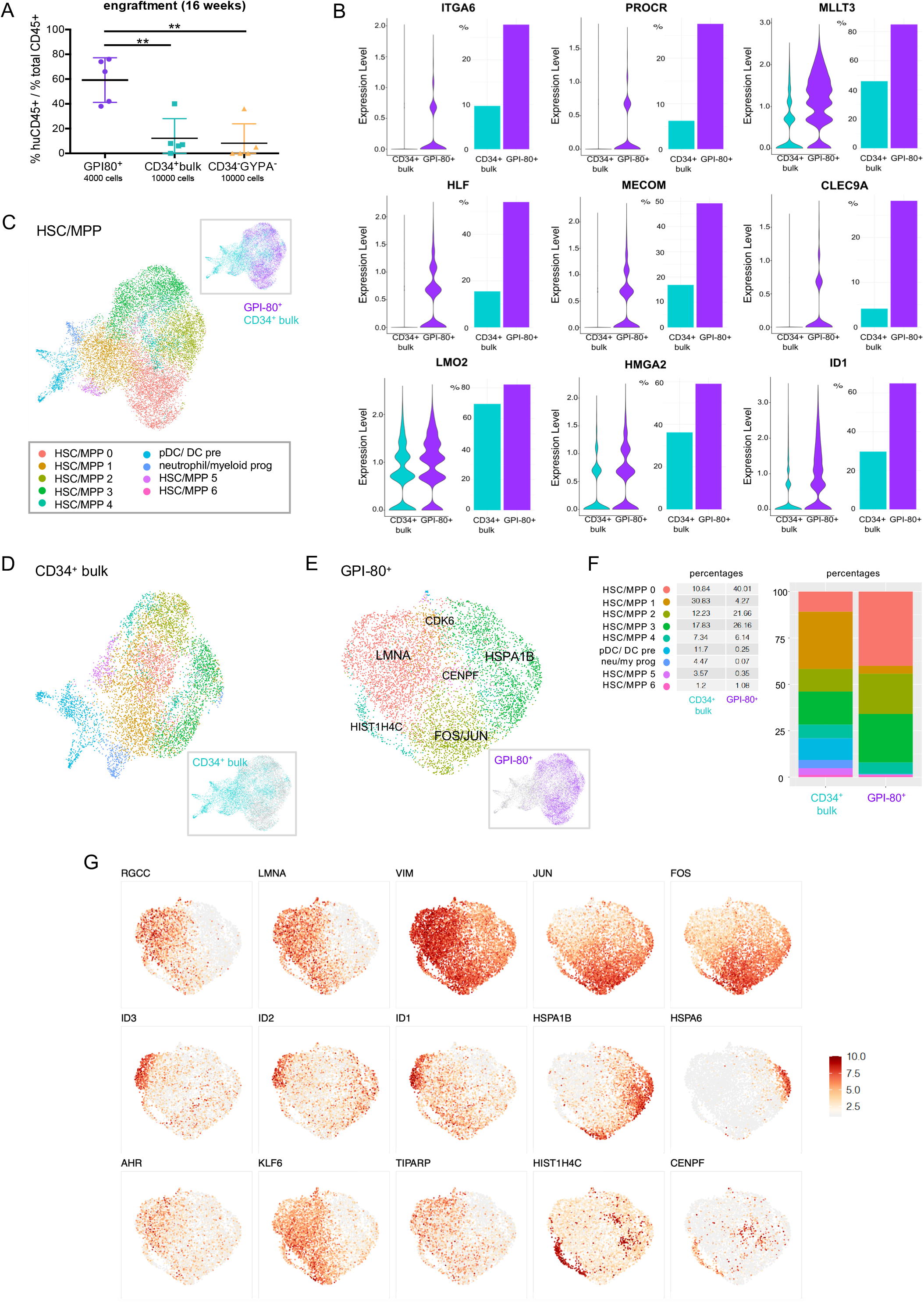
Single cell transcriptomic profiling of the engraftment potential of FL-derived HSCs. (A) Engraftment data demonstrating superior engraftment of the GPI-80^+^ fraction 16 weeks post transplantation. Five NSG mice were transplanted per condition with 10,000 cells each (4,000 cells for the GPI-80+ condition) and engraftment was assessed by determining the ratio (%) of peripheral blood cells expressing human CD45 versus total CD45 expressing cells. The mean is shown with error bars representing the standard error of the mean. *\*p < 0.05, **p < 0.01* (B) Violin plots of differentially expressed genes between HSCs/MPPs in the GPI-80^+^ and CD34^+^ bulk fractions (left) and bar plots illustrating the percentage of cells expressing these genes per sample (right). (C) UMAP representation showing the clusters identified in the CD34^+^ HSC/MPP population. (D) UMAP representation of the HSCs/MPPs corresponding to the CD34^+^ bulk sample, retaining the cluster definitions established in panel C. (E) UMAP representation of the HSCs/MPPs corresponding to the GPI-80^+^ sample, retaining the cluster definitions established in panel C. Additional labeling with one of the top most enriched genes is shown for each cluster (see **Table S4**). (F) Cluster dynamics upon GPI-80-enrichment showing the proportion of each cluster before (CD34^+^bulk) and after (GPI-80^+^) functional enrichment in a stacked bar plot. Percentages of cells in each cluster are shown on the left. (G) UMAP overlays illustrating mRNA expression patterns of genes enriched in different GPI-80^+^ clusters. The scale bar shows scaled expression from 0-Max.

In concordance with their superior engraftment potential, we found enrichment for known HSC markers such as *ITGA6* (*CD49f*), *PROCR* (*EPCR* or *CD201*), *MLLT3*, *HLF*, *MECOM*, *CLEC9A*, *LMO2* and *HMGA2* in the GPI-80^+^ fraction compared to CD34^+^bulk cells (**Figure 2B**). Enrichment for these genes, and others with currently unexplored connections to HSCs, distinguishes the GPI-80^+^ fraction from bulk CD34^+^ cells, revealing for the first time, a detailed transcriptomic signature that marks engraftable FL HSCs. A comprehensive list of all differentially expressed genes (DEGs) that make up this engraftment signature is presented in **Table S2**.

To further analyze engraftable FL HSCs, we first focused on the CD34^+^ HSC/MPP population (**Figure 1C**) captured within the grey dotted circle in **Figure 1A**. Separate clustering of this subset resulted in seven distinct HSC/MPP clusters (HSC/MPP 0-6) in addition to previously identified neutrophil/myeloid progenitor and pDC/DCpre clusters (**Figure 2C**). To determine which of these clusters best correlated with engraftment potential, we tracked cluster dynamics upon GPI-80^+^ enrichment, which we showed was associated with enrichment for functional HSCs. Using the previously determined cluster designations from **Figure 2C**, we identified the corresponding clusters in the individual samples making up this CD34^+^ HSC/MPP fraction (**Figures 2D-E**). Mapping the changes in cluster proportions between CD34^+^ bulk and GPI-80^+^ enriched samples, showed a reduction of certain clusters with an increase of others (**Figure 2F**). The relative absence of cells corresponding to HSC/MPP cluster 1 within the GPI-80+ enriched sample suggests that these cells and their associated transcriptomic signature likely contribute minimally to the engraftment potential of FL HSCs. Emboldening this point, one of the top enriched genes in this cluster is *CDK6* which has been described as an activation marker, marking the transition towards more differentiated progenitors (Laurenti et al., 2015). Similarly, we noted a significant reduction in both cluster 5 and those clusters corresponding to pDC/ DCpre cells and neutrophil/myeloid progenitors. Conversely, the proportion of cells in clusters 0, 2 and 3 increased in the GPI-80^+^ sample compared to the CD34^+^ bulk population. Of these, the most pronounced change was noted for HSC/MPP cluster 0, which showed a substantial 4-fold increase. This specific enrichment, resulting in cluster 0 cells accounting for 40% of the total GPI-80^+^ population, strongly suggests that the transcriptomic profile corresponding to this cluster best represents cells with functional engraftment potential. The proportions of HSC/MPP clusters 4 and 6, representing cells engaged in replication, remained stable upon GPI-80^+^ enrichment (**Figure 2F**, **Figure S2A**). Overall these cluster dynamics resulted in a slight increase in cells in G1 (or G0) phase at the expense of cells in S/G2/M phase in the GPI-80^+^ fraction compared to the CD34^+^bulk fraction (**Figure S2B**).

To further dissect the gene expression signature specific to cluster 0, which we found preferentially enriched within the GPI-80^+^ sample, we assessed the differential gene expression between this and other GPI-80^+^ clusters. *RGCC* and *LMNA* were identified as the top enriched genes in cluster 0, closely followed by *VIM* and *ID1* and *ID3* (**Figure 2G**, **Table S3, Figure S2C**). Interestingly, *LMNA* was found expressed at higher levels in FL than in postnatal CD34^+^ cells (**Figure S2D**) in line with its previously described inverse correlation with age (Adelman et al., 2019). In addition to ID signaling pathway members, whose expression appears concentrated along the outer edge of cluster 0, we also found enrichment for *AHR, KLF6* and *TIPARP,* key players in the aryl hydrocarbon receptor (AHR) pathway (**Figure 2G**, **Table S3**).

Although less prominent than what we observed for cluster 0, the proportion of cells in clusters 2 and 3 also increased upon GPI-80 enrichment. Of these, cluster 2 was characterized by enrichment for *FOS* and *JUN* expression, two important factors involved in numerous signaling pathways (**Figure 2G**, **Table S3**, **Figure S3E**). Interestingly, cluster 3 was enriched for heat shock proteins (*HSPA1A*, *HSPB1B*, *HSPA6*, *HSPB1*), suggesting that the unfolded protein response best distinguishes this population (**Figure S3F**). Together, these signatures suggest that the GPI-80^+^ fraction harbors several subtypes of HSCs with different characteristics. However, the dominance of enrichment by GPI-80 makes a strong case for the cluster 0 expression profile to be the most likely representation of engraftable FL HSCs.

### Antibody-derived tags (ADTs) link cell surface protein expression to transcriptomic characterization of FL HSCs

Sequencing of antibody-derived tags (ADTs) via CITE-seq enabled the coupling of transcriptional data with cell surface marker expression. Overall, a strong correlation was found between mRNA and ADT expression, with the latter providing increased coverage due to lower dropout rates (Stoeckius et al., 2017) (**Figure 3A-B**, **Figure S3A-B**). **Figure 3A** projects both layers of information onto the CD34^+^bulk UMAP for a selection of HSC markers. Notably, for CD34, CD90 and CD49f, often used in combination to enrich for HSCs, ADT expression patterns were observed that were not readily apparent based on mRNA data alone, indicating co-expression of these markers at the protein level. Another observation that was evident from the analysis of ADT but not mRNA expression data, was enrichment for CD201 expression in cluster 0 both in the CD34^+^bulk and GPI-80^+^ fractions (**Figure 3A-B, Figure S3B, Table S3**). When comparing cell surface markers included in our ADT panel for their ability to identify cluster 0, CD201 stood out. This was evident for both CD34^+^ HSC/MPP fractions (**Figure 3C-D**) and also throughout the 26,407 combined cells from all 4 original samples, irrespective of CD34 enrichment (**Figure 3E**). This suggests that CD201 could be harnessed as a single marker to specifically enrich for highly functional FL HSCs. To this point, progressive *in silico* enrichment for CD201 expression demonstrated a robust increase in the proportion of cells corresponding to the engraftment signature represented by cluster 0 (**Figure 3F**). Notably, 49% of the top CD201 expressing cells within the CD34^+^ bulk fraction represented cluster 0 cells versus ~25% of cells when considering markers such as CD49f, CD90 and CD133 (**Figure 3C,F**). This level of enrichment for cluster 0 even surpassed that observed when using GPI-80 to sort functional HSCs from bulk CD34^+^ cells (**Figure 2F**). In stark contrast to CD201, enrichment based on surface expression of CD164 was correlated with a decrease in cluster 0 frequency (**Figure 3G**). Furthermore, progressive CD34 enrichment resulted in a moderate reduction in cluster 0 combined with reductions in clusters 2 and 3 and an increase in cluster 4 (**Figure 3H)**. Progressive enrichment for other HSC markers such as CD90, CD49f and ENG (endoglin, CD105) did not alter the transcriptomic profile of cells (**Figure S3C**).

**Figure 3.**
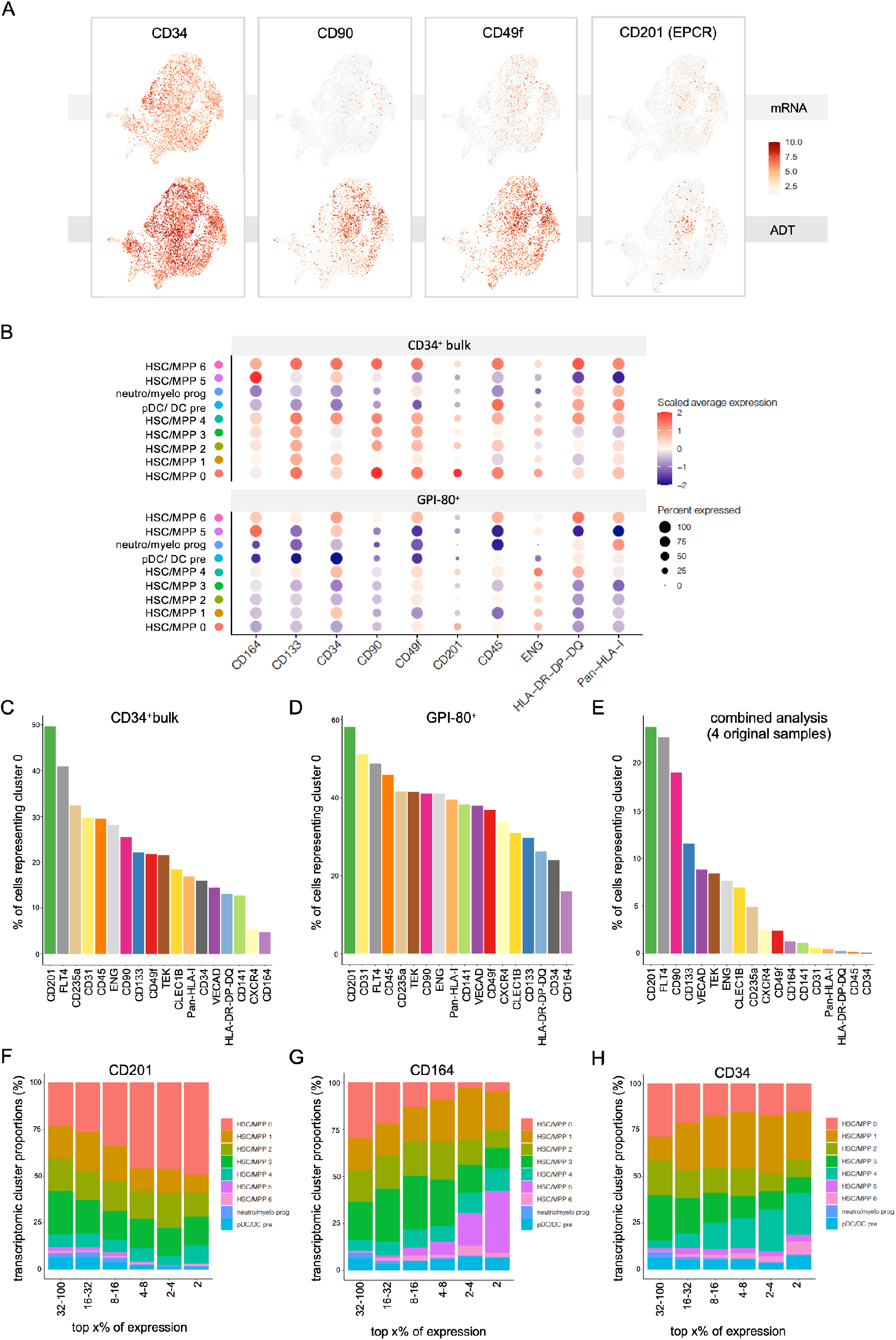
Antibody-derived tags (ADTs) link cell surface protein expression to transcriptomic characterization of FL HSCs. (A) Comparison of mRNA and ADT expression for a selection of HSC markers in the CD34^+^bulk sample. (B) Dot plot illustrating the distribution of antibody-derived tag (ADT) expression for a selection of HSC markers in the GPI-80^+^ sample versus the CD34^+^ bulk sample. Expression is represented as z-scores and the scale is kept constant between the CD34^+^ bulk and GPI-80^+^ sample for each marker to allow for comparison of relative intensities. “Percent expressed” represents the fraction of cells having non-zero expression values for each cell cluster in the two samples. (C-E) Comparison of the ability to enrich for cluster 0 across the different cell surface markers included in our ADT panel for the CD34^+^bulk fraction (C), GPI-80^+^ fraction (D) and combined analysis of all 4 originally sorted samples (E). The proportion of cells representing cluster 0 (%) is shown on the y-axis, considering cells enriched for each marker (top 5%) on the x-axis. (F-H) Transcriptomic cluster composition (%) within the CD34^+^ bulk fraction upon stepwise enrichment for cell surface marker expression of CD201 (F), CD164 (G) and CD34 (H). The x-axis is divided in segments showing the indicated top x% of expression for each marker.

Together, these findings illustrate that cell surface protein expression coupled with transcript-level data provides a powerful approach towards classifying potential enrichment strategies for highly functional HSCs. It also provides a complimentary, yet orthogonal methodology with which to tease apart the heterogeneity that exists within the HSC/MPP compartment of the FL.

### Multi-dimensional flow cytometric characterization across multiple FL samples

To further extend the cell surface marker characterization of FL HSPCs, five additional CD34^+^ FL samples (week 16-22) were profiled by flow cytometry using a 21-marker antibody panel (**Table S5**). Flow cytometry, considered the gold standard for assessing cell surface protein expression, provides an alternative approach to validate ADT-based CITE-Seq and allows for the profiling of very large numbers of cells. To enable comprehensive analysis of the resulting multi-dimensional single cell protein expression dataset, marker expression was projected onto a common UMAP scaffold analogous to representations of high-dimensional transcriptomic data. Healthy adult peripheral blood mononuclear cells (PBMCs) were simultaneously phenotyped as a control and projected into the same UMAP space to provide context for less mature FL cell subsets and Phenograph clustering (Levine et al., 2015) was performed on the combined data (PBMCs and FL CD34^+^ cells) (**Figure 4A-D, Figure S4A-B**). While some variability was noted between FL samples, each of the identified FL clusters was populated with cells from more than one FL sample (**Figure S4A,C**). As expected, certain Phenograph clusters were only represented in the PBMC sample as they contained the more mature cell populations. In addition to HSC markers present in our CITE-seq panel, lineage markers were also included to identify committed progenitors within the CD34^+^ FL fraction. UMAP visualization of the combined flow data from 5 CD34^+^ FL samples allowed for assessment of the overlap between CD34 expression and that of negative selection markers such as CD38 and CD45RA, as well as more committed lineage markers (**Figure 4E**). This process allowed for clear sub-fractionation of the mature PBMCs (**Figure 4D-E**) and demonstrated that certain mature cell markers (CD66c, CD33) are also broadly expressed within the CD34^+^ population, highlighting the importance of carefully choosing such markers in negative selection strategies (Taussig et al., 2005). Notably, when looking at the portion of the UMAP that represents CD34^+^CD38^−^CD45RA^−^ cells (clusters 1, 3, 8, 13, 30, 31 in **Figure 4D**), cells that also co-express CD90 and CD49f (right edge of clusters 1 and 3) were identified as suggested by our ADT data (**Figure 3A**). In this CD34^+^CD38^−^ CD45RA^−^ fraction, expression of other established HSC markers such as CD201 and GPI-80 was evident. Interestingly, this visualization drew attention to a population of cells at the tip of cluster 3 in which CD49f, CD201 and GPI-80 appeared to be co-expressed (**Figure 4D-E**). Protein-level co-expression of these HSC markers could be inferred based on comparison of ADT expression patterns in the CD34^+^bulk and GPI-80^+^ fractions as well (**Figure 3A-B, Figure S3B**). Taken together, this multi-dimensional flow characterization confirms the expression patterns observed with ADTs and validates these findings across biologically distinct FL samples.

**Figure 4.**
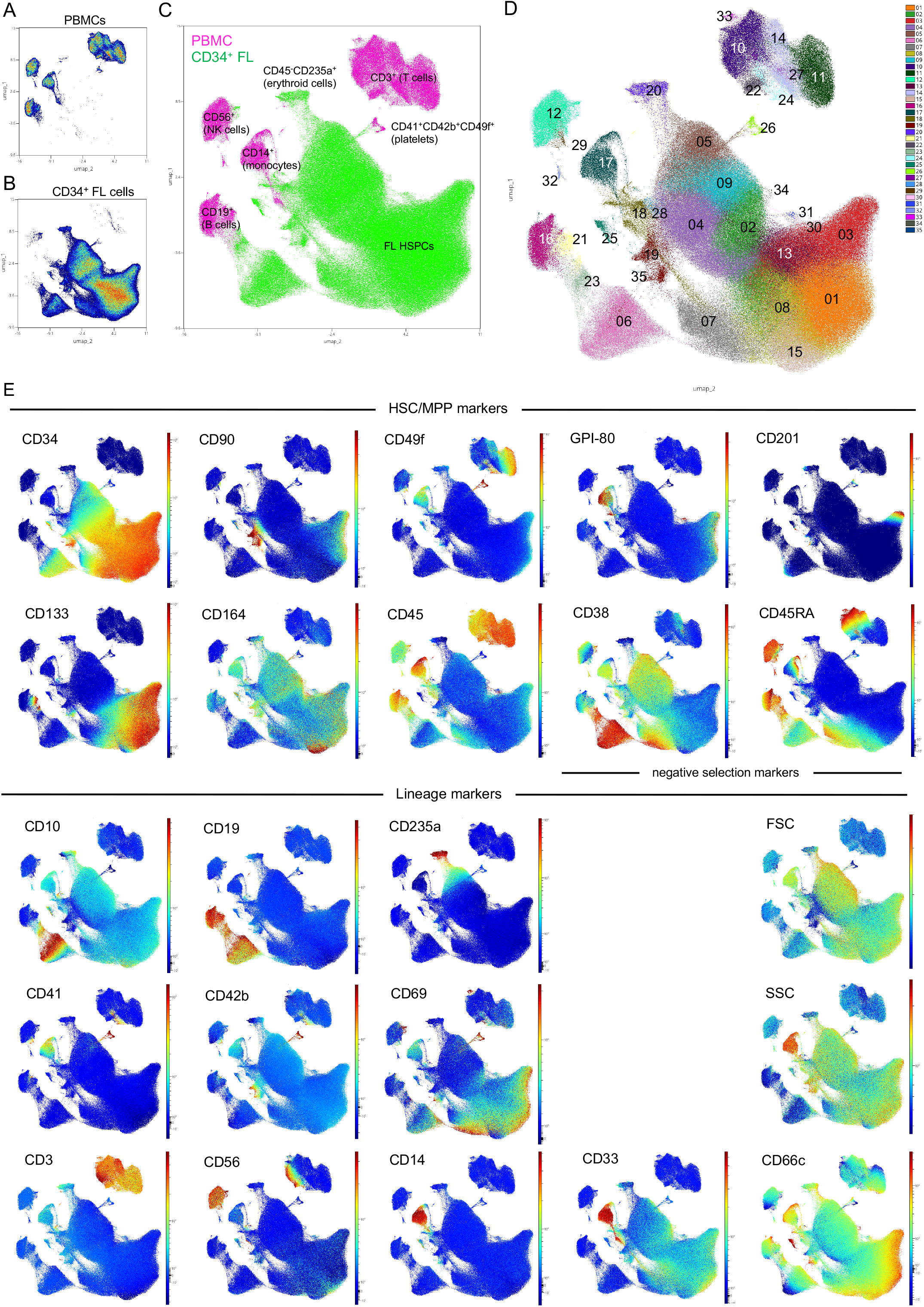
Multi-dimensional flow cytometric characterization across multiple FL samples. (A-B) Individual representation of PBMCs (A) and CD34^+^ FL cells (B) in the same UMAP space. Color indicates plot density (C) UMAP representation of both PBMCs (pink) and CD34^+^FL cells (green). Major mature blood cell subsets are indicated on the PBMC sample for orientation purposes. (D) UMAP representing the different Phenograph clusters identified in the combined dataset (PBMCs and CD34^+^ FL cells). (E) Expression patterns of individual markers and light scatter cytometric parameters overlaid onto the UMAP representation of the combined data. Color indicates fluorescence or light scatter signal intensity.

### Integrating gene expression and cell surface marker information to visualize and compare HSC enrichment strategies

Inspired by the co-expression pattern of classical HSC markers CD34, CD90 and CD49f highlighted by ADTs (**Figure 3A**) and confirmed by flow data (**Figure 4E**), we asked whether we could use both levels of information (mRNA and ADT expression) to visualize and compare HSC enrichment strategies via *in silico* sorting. This bioinformatic approach harnesses ADT expression values to recreate gating strategies used in FACS sorting. Three sorting strategies were compared by this method: 1) The ‘classical’ HSC enrichment scheme (lin^−^ CD34^+^CD38^−^CD45RA^−^CD90^+^CD49^+^) (Notta et al., 2011); 2) an HSC enrichment strategy that was recently described to yield a highly purified population of engraftable HSCs from CB (lin^−^CD34^+^CD38^−^CD133^+^GPI-80^+^), which we refer to as the ‘Sumide et al.’ signature (Sumide et al., 2018); and 3) an ‘EPCR^+^’ signature (CD34^+^CD38^−^CD201^+^). Endothelial protein C receptor (EPCR, also known as PROCR or CD201) has been described in mouse and humans to mark the LT-HSC and appears to be stably retained on the cell surface when culturing HSCs *ex vivo* (Balazs et al., 2006; Fares et al., 2017; Subramaniam et al., 2019). After *in silico* sorting of cells corresponding to these signatures, the outcomes of the three HSC enrichment strategies were visualized via projection onto a common UMAP reference frame and their transcriptomic profiles were further compared (**Figure 5A-C**). An overview of the *in silico* sorting strategy is presented in **Figure S5A**, whereby CD34^+^CD38^−^ cells were gated and then sub-gated via additional positive selection markers of each signature (described in detail in methods section). This process was guided by our flow cytometry experiments to ensure that the *in silico* gating reflected populations that would be obtained via analogous FACS sorting (**Figure S5A-B**). Overlaying the corresponding cells onto the CD34^+^bulk UMAP suggested that considerable similarity exists between the sort strategies, with all three approaches enriching for cluster 0 cells (**Figure 5A**). The majority of DEGs for both the classical (82.94%) and EPCR^+^ (80.12%) signatures were shared with the Sumide et al. signature (**Figure 5B),** which closely matches our GPI-80-based functional enrichment strategy. The strong correlation between the top enriched genes further corroborates the extensive transcriptomic overlap between cells captured by these different HSC enrichment strategies (**Figure 5C**). Together these data suggest that the transcriptomic signature of the engraftable HSC identified in this work is not exclusive to the GPI-80^+^ enrichment strategy, but rather represents a generalizable engraftment signature for FL HSCs irrespective of purification strategy.

**Figure 5.**
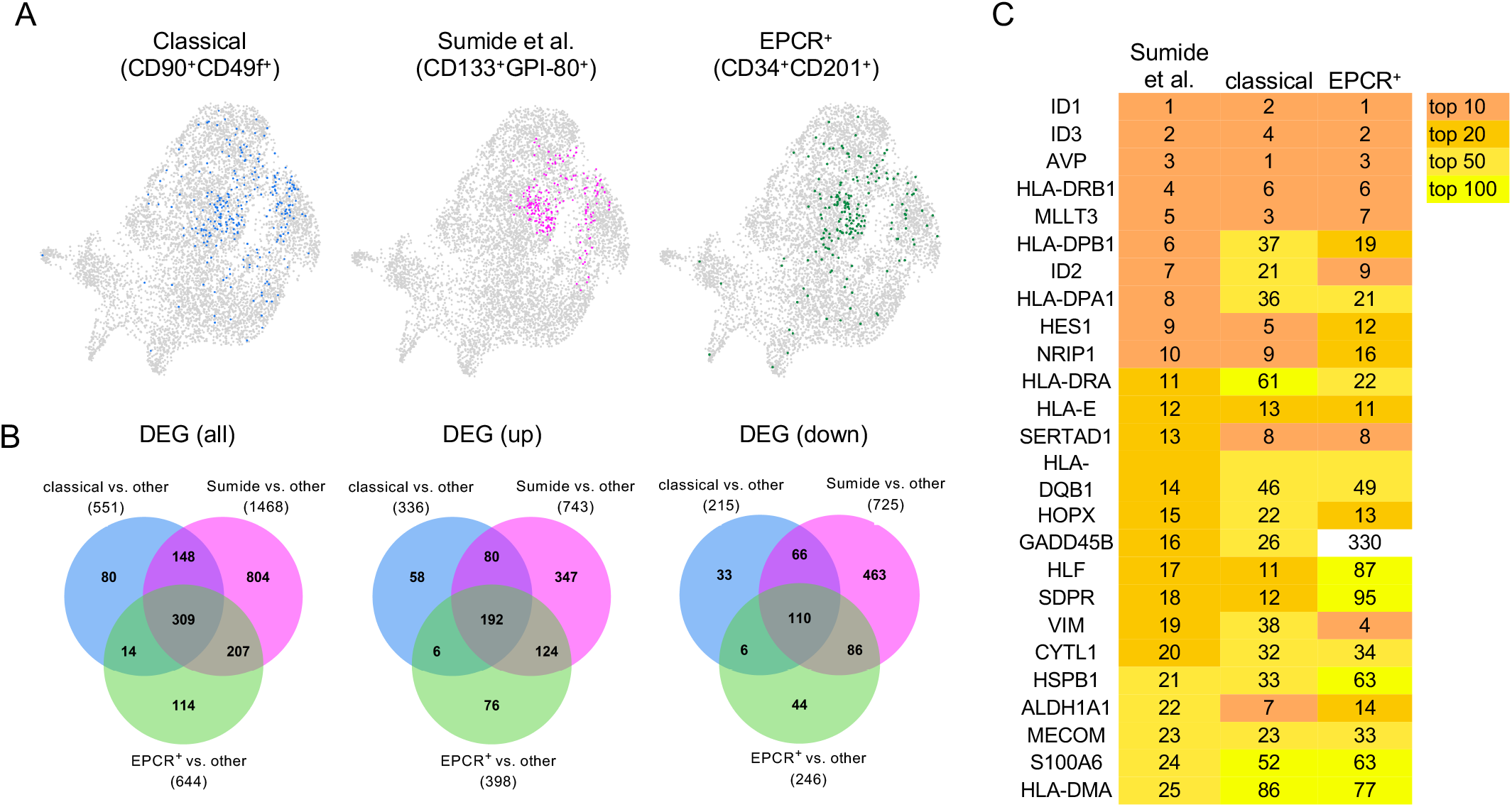
Integrating cell surface marker and gene expression information to visualize and compare HSC enrichment signatures. (A) Overlays of in silico sorted HSC enrichment signatures onto the CD34^+^bulk UMAP (B) Venn diagrams showing overlap of differentially expressed genes (DEGs) between the different HSC enrichment signatures. For each signature, the number of DEGs between cells corresponding to that signature based on *in silico* sorting and the background (other) is depicted. (C) Overview of top 25 enriched genes corresponding to the Sumide et al. signature and their rank in the different HSC enrichment signatures.

## DISCUSSION

Our examination of the molecular programs associated with potent engraftment capacity can serve as a resource which we hope will empower a broad range of future studies. From the development of strategies that preserve HSC functionality ex vivo, to illuminating the developmental roadmap for the generation of HSCs from PSCs, our data reveals a clear transcriptional and protein-level program against which to benchmark these approaches.

To arrive at an in-depth characterization of engraftable FL HSCs, we harnessed the functional HSC enrichment capacity of GPI-80, a highly effective marker associated with engraftment (Prashad et al., 2015; Sumide et al., 2018). Despite representing only 2.37% of total CD34^+^ cells, sorting thousands of CD34^+^GPI-80^+^ cells allowed us to perform parallel profiling of this rare population at a functional, transcriptional and surface protein level. After confirming their superior engraftment potential via xeno-transplantation, we mined the transcriptome of these rare cells, resulting in a molecular signature of engraftable FL HSCs at unprecedented resolution.

Given the transcriptionally distinct clusters we discerned within this highly purified population, we sought to understand which of these transcriptomic profiles exhibited the strongest correlation with engraftment potential. By analyzing cluster proportions before and after enrichment, we identified three clusters that were significantly enriched, thus corresponding to putative engraftable HSCs.

The first cluster (HSC/MPP; cluster 3) was characterized by a prominent unfolded protein response (UPR) signature, represented by an abundance of heat shock proteins (*HSPA1A*, *HSPB1B*, *HSPA6*, *HSPB1*). Interestingly, protein quality control has been linked to the ability of HSCs to maintain their undifferentiated status (Hidalgo San Jose et al., 2020; van Galen et al., 2014a). FL HSCs possess the unique capacity to tolerate significant proliferation without impacting multilineage engraftment potential and this has been associated with a heightened DNA damage response (DDR) in FL versus postnatal HSCs (Biswas et al., 2020). Similarly, there appears to be a role for the maintenance of proteome integrity in shielding HSC function during expansion in the FL. While bile acids have been described to serve as chaperones alleviating unfolded protein stress in expanding mouse FL HSCs (Sigurdsson et al., 2016), our data suggest that heat shock proteins may take on this role in the human FL.

The second cluster (HSC/MPP; cluster 2) over-represented in highly enriched FL HSCs exhibited a series of upregulated signaling pathways. Amongst these, a variety of interleukin pathways were represented, suggesting that these cells may have altered responsiveness to interleukin signaling. The most enriched genes were *FOS* and *JUN*, two downstream effectors of the JNK pathway, which has recently been shown to play a critical role in regulating HSC expansion (Xiao et al., 2019).

The third and most highly-represented cluster upon functional enrichment (HSC/MPP cluster 0) exhibited increased expression of *RGCC, LMNA,* ID genes and members of the AHR pathway (*AHR, KLF6, TIPARP and JUND*). AHR modulators such as SR-1 have been described to expand CD34^+^ CB cells (Boitano et al., 2010) and we and others have linked AHR inhibition to enhanced endothelial-to-hematopoietic transition and HSPC expansion during in vitro differentiation from PSCs (Angelos et al., 2017; Leung et al., 2018), highlighting a role for the AHR pathway in HSPC biology. *LMNA* was identified as the second most enriched gene in this cluster. *LMNA* encodes the nuclear lamina protein Lamin A/C and is expressed in postnatal HSCs where it shows a decline in expression throughout aging (Adelman et al., 2019; Grigoryan et al., 2018). Interestingly, comparing *LMNA* expression between our FL CD34^+^ bulk cells and existing scRNAseq data sets representing postnatal CD34^+^ fractions (Belluschi et al., 2018; Velten et al., 2017; Zheng et al., 2018), revealed that FL CD34^+^ cells possessed the highest *LMNA* expression (**Figure S3D**), suggesting that the decline in *LMNA* expression might begin in utero. Given the superior engraftment potential of FL HSCs compared to postnatal HSCs (Holyoake et al., 1999), this might suggest a role for *LMNA* in endowing FL HSCs with this remarkable potential. Consistent with this hypothesis, *LMNA* expression has been described to be more prominently expressed in LT-HSCs than in less potent ST-HSCs in mice (Grigoryan et al., 2018), a finding that is reflected in our data by the enrichment of *LMNA* in the GPI-80^+^ compared to the CD34^+^ bulk fraction (**Table S2**). Together these findings support a link between *LMNA* expression and HSC functionality.

We also observed enrichment for members of the inhibitor of DNA binding (ID) gene family along the top edge of cluster 0. Expression of *ID1*, *ID2* and *ID3* has been reported to be higher in HSCs as compared to downstream hematopoietic progenitors in CB (van Galen et al., 2014b). Furthermore, reduction in the expression of these genes has been shown to coincide with loss of quiescence and *in vivo* repopulating capacity (van Galen et al., 2014b). Moreover, ID1 has recently been implicated in controlling the balance between dividing and resting neural stem cells by promoting quiescence (Zhang et al., 2020). It is tempting to speculate that ID genes could play a similar role in FL HSCs, especially when taking into consideration co-enrichment for other factors involved in cell cycle control in this cluster such as *RGCC* or ‘regulator of cell cycle’. Overall, the preferential enrichment of cluster 0 upon functional enrichment, coupled with the increases in ID gene expression, suggest that the transcriptomic profile of cluster 0 most likely represents engraftable FL HSCs.

Notably, a recent study profiling PSC-derived HSPCs at the single cell level, identified *ID2* as enriched in what was considered the most naïve in vitro-derived HSPC fraction, highlighting an encouraging overlap in gene expression with engraftable HSCs in vivo (Fidanza et al., 2020). In this work, the transcriptomic profile of PSC-derived HSPCs was compared to that from FL HSPCs using an artificial neural network, identifying additional commonalities but also dissimilarities that might suggest how to further improve PSC-based HSC generation. In line with this goal, we believe that our comprehensive characterization of engraftable HSCs and the detailed engraftment signature that cluster 0 represents will be of particular interest to the field to further optimize this process.

Beyond in-depth transcriptional characterization, this study also provides the first linked cell surface marker expression data of engraftable FL HSCs. Comparing both transcript and protein expression data, we found that ADT read-outs offered complementary insights as they allowed for the identification of marker co-expression that was not readily apparent based on mRNA expression alone. Moreover, ADT but not mRNA expression data suggested that EPCR (CD201) could be used as a cell surface marker to specifically enrich for cluster 0 cells and thus engraftable FL HSCs. While this marker has been proposed as a useful addition to existing enrichment strategies to purify FL HSCs (Subramaniam et al., 2019), the linked mRNA and ADT data in this work would suggest that EPCR may serve as a viable single enrichment marker for functional FL HSCs.

Using flow cytometry, we further extended our protein level characterization of FL HSPCs through simultaneous assessment of 21 cell surface markers and confirmed the expression patterns observed based on ADT expression data as representative across multiple biological replicates of FL. The UMAP representation of the multi-dimensional flow data provides a comprehensive overview of expression patterns of both HSPC and lineage markers and thus allows for a straightforward visual comparison of different HSC enrichment strategies. This visualization highlighted a region characterized by co-expression of several prominent HSC enrichment markers, suggesting that it could represent the apex of the hematopoietic hierarchy where engraftable HSCs reside. This interpretation was further supported by the almost exclusive localization of CD201 expression in this region as our ADT data suggested that CD201 expression marks cluster 0 cells highly enriched in engraftment potential. These observations are in line with the recent molecular characterization of mouse HSCs where *Procr* (*Epcr*) was shown to be enriched in the most primitive subset of functional long-term repopulating HSCs (Rodriguez-Fraticelli et al., 2020). Our flow cytometry data set can be accessed and further interrogated via the following repository (http://flowrepository.org/id/FR-FCM-Z32M).

Lastly, to illustrate the added value of cell surface marker expression data linked to transcriptomic data we integrated ADT and mRNA information to compare three well-established HSC enrichment strategies. Through the process of *in silico* sorting, we gated populations of interest based on ADT expression values and compared the transcriptomic profiles of the resulting cells. This analysis indicated that a GPI-80-based enrichment strategy showed very high similarity to enrichment strategies driven by either CD90^+^CD49f^+^ or EPCR^+^ enrichment. This led us to conclude that the engraftment signature that we identify in this work is representative of engraftable FL HSCs irrespective of the markers used during isolation. Using our dataset, the same *in silico* sorting approach can also be applied to find answers to different queries without the need to physically sort out cell fractions and subject them to transcriptomic profiling.

The engraftment signature as well as the ‘refined engraftment signature’ resulting from further dissection of the transcriptional profiles present in the GPI-80^+^ enriched fraction are available in their entirety (**Table S2-3**) and the CITE-seq data set described in this work has been made available in an interactive format at https://engraftable-hsc.cells.ucsc.edu. We envision that this transcriptional and protein-level profiling of engraftable FL HSCs will serve as an openly shared resource, enabling new biological insights into engraftment potential, including how it can be retained during *ex vivo* culture and how it can be induced to generate functional HSCs from PSCs.

### Limitations of study

This study presents the in-depth characterization of engraftment potential at a single time point in hematopoietic development where we have single cell profiled 26,407 cells, specifically focusing on a population highly enriched in engraftable FL HSCs. Although the resulting transcriptomic and linked ADT expression data reflect a single time point, we have verified that the identified cell surface marker expression patterns are representative of FL HSPCs via multiparameter flow cytometry of five distinct FL samples ranging from 16-22 weeks of developmental age. Moreover, we found striking similarities between the hematopoietic landscape represented by our combined data set and that reported in earlier work (Popescu et al 2019). Further supporting our findings are the presence of genes previously identified to be enriched in FL HSPCs fractions such as *MLLT3*, *CLEC9A*, and *SPINK2* (Calvanese et al., 2019; Popescu et al., 2019). Importantly, the unprecedented resolution at which the engraftable HSC fraction is interrogated in this study, together with the multi-modal profiling of these cells at the functional, transcriptomic and protein-level, makes our in-depth characterization and resulting engraftment signature unique among prior studies interrogating HSPCs.

## ACKNOWLEDGEMENTS

The authors would like to thank Dr. Yuriy Alekseyev and Ashley Leclerc from the Boston University School of medicine (BUSM) Microarray and Sequencing Resource Core Facility for their assistance as well as Brian Tilton from the BUSM Flow Cytometry Core Facility. The mouse cells that were mixed in as background controls were kindly provided by Dr. Sara Lewandowski and Dr. Laertis Ikonomou.

## AUTHOR CONTRIBUTIONS

Conceptualization, K.V., G.J.M. and A.B.B.; Methodology, K.V., R.D., J.D.C., A.C.B., G.J.M. and A.B.B.; Software, C.V.-M., J.L.-V., Z.W., R.D., A.C.B.; Formal Analysis, C.V.-M., J.L.-V., Z.W., T.M.M., M.L., J.D.C., R.D., A.C.B.; Investigation, K.V., W.F.G.-B., V.V., T.W.D., S.K., A.C.B.; Writing – Original Draft, K.V., G.J.M. and A.B.B.; Writing – Review & Editing, K.V., G.J.M. and A.B.B.; Data curation, C.V.-M., A.C.B.; Supervision, K.V., G.J.M. and A.B.B.

## METHODS

### RESOURCE AVAILABILITY

#### Lead Contact

Further information and requests for resources and reagents should be directed to and will be fulfilled by the Lead Contact, George J Murphy

#### Materials Availability

This study did not generate new unique reagents.

#### Data and Code Availability

The datasets/code generated during this study are publicly available at:

CITE-seq dataset/code (raw data): GEO accession number: GSE160251 https://www.ncbi.nlm.nih.gov/geo/query/acc.cgi?acc=GSE160251

CITE-seq dataset/code (interactive format): https://engraftable-hsc.cells.ucsc.edu

Multiparameter flow cytometry data/code: Unmixed fluorescence cytometry dataset with integrated UMAP and Phenograph parameters is available at http://flowrepository.org/id/FR-FCM-Z32M

### EXPERIMENTAL MODEL AND SUBJECT DETAILS

#### Ethics Statement

All mouse research complied with the Institutional Animal Care and Use Committee (IACUC) of the Massachusetts General Hospital (Protocol #2009N000136) and use of human tissues was approved by Partners Human Research Committee (Protocol #2016P001106).

#### Fetal liver samples

The stage of the fetal liver samples is indicated as developmental age (two weeks less than the gestational age, which is counted from the last menstrual period). The fetal liver sample used for the CITE-seq analysis was collected at 21 weeks (sex unknown). For the multiparameter flow characterization the following samples were used: FL1 (sex unknown, 21 weeks), FL2 (F, 22 weeks), FL3 (sex unknown, 16 weeks), FL4 (M, 17 weeks), FL5 (F, 17 weeks).

#### Peripheral blood mononuclear cells (PBMCs)

PBMCs from a 65 year old individual (M) were used as a control for the flow cytometry experiments (New York Biologics Inc).

#### Mouse embryonic stem cells (mESCs)

mESCs from Nkx2-1^mCherry^ mice (Bilodeau et al., 2014) were spiked into samples prepped for CITE-Seq to control for background staining from the TotalSeq A antibodies. Undifferentiated mESCs were maintained on a feeder layer of mitotically inactivated mouse embryonic fibroblasts (MEFs) in serum-containing media consisting of DMEM (Life Technologies, 11995-073) with 15% FBS (Thermo Fisher Scientific, NC0712155), 200 mM L-glutamine (Invitrogen, 25030-164) and 100 μg/ml Primocin (Thermo Fisher Scientific, NC9392943), supplemented with LIF-containing conditioned media (Millipore, ESG1106) at 1 U/ml. mESCs were passaged as needed when cultures reached appropriate confluency using 0.05% trypsin (Invitrogen, 25300-120)

#### *NOD-*scid IL2rγ^null^*(NSG) mice*

NOD-*scid IL2rγ^null^* (NSG) mice (F, 14-16 weeks old) were used in transplantation experiments.

### METHOD DETAILS

#### Processing of FL samples

FL samples were mechanically dissociated into small pieces and subsequently incubated in Liver Digest Medium (Fisher Scientific, 17703034) at 37°C. Mononuclear cells were isolated over a Ficoll gradient (Lymphoprep: Stem Cell Technologies, 7851) prior to separation into CD34^+^ and flowthrough (CD34^−^) cells using magnetic beads (CD34 Microbead Kit: Miltenyi Biotec, 130-046-702).

#### CITE-seq sample preparation

FL cells were thawed and allowed to recover at 37°C for an hour prior to staining. Cells were blocked with TruStain FcX (BioLegend, 422301) and stained with TotalSeq A antibody mix containing 1ug of each TotalSeq A antibody per sample (BioLegend, see **Table S1** for a list of antibodies). Anti-human CD235a-APC antibody (BD Biosciences, 551336) was added to the CD34^−^ sample. The CD34^+^ fraction was stained with anti-human CD34-APC antibody (BD Biosciences, 555824) and anti-human GPI-80-PE antibody (MBL International, D087-5). All samples were stained with calcein blue (Invitrogen, C34853) for live/dead exclusion and cell populations were sorted as shown in Figure S1A prior to loading onto the 10X Genomics platform. Chromium Single Cell 3**ʹ** Reagent Kit v3 with Feature Barcoding technology for Cell Surface Protein was used and the recommendations from the manufacturer were followed as specified in 10x Genomics user guide document number CG000185, Rev B.

#### Transplantation experiments

Cells from the same sorted fractions as used in the CITE-seq experiment were used for transplantation into NOD-*scid IL2rγ*^*null*^ (NSG) mice. Five mice were transplanted per condition with 10,000 cells each (4,000 cells for the GPI-80+ condition) and engraftment was assessed after 16 weeks by determining the ratio (%) of peripheral blood cells expressing human CD45 versus total CD45 expressing cells. Cells were stained with APC-conjugated anti-human CD45 antibody (BioLegend, 304012) and anti-mouse CD45-PacBlue antibody (BioLegend, 103126).

#### CITE-seq: transcriptomic analysis

Fastq files were generated and counts extracted from each of the three runs separately for expression libraries and antibody-derived tag (ADT) libraries using bcl2fastq v.2.2 and cellranger v.3.0.2. The expression libraries were mapped to a combination of the human and mouse genome references (GRCh38 and GRCm38), in order to detect both human and mouse cells. The ADT counts were summarized using CITE-Seq-Count v 1.4.2. We used Seurat v.3 to further process the data. The proportion of cells which included both human and mouse genes, was used as a threshold to estimate the empirical doublet rate. As expected based on the 10X Chromium guidelines, the doublet rate was proportional to the density of cell loading, but was found to be half the rate originally calculated. Cells with more than a 25% of reads mapping to mitochondrial genes were filtered out. The transcriptomic and ADT assays were aggregated into an integrated multi-assay analysis. The CD34^−^GYPA^−^ (CD34N-CD235AN) sample generated 10,253 cells at a depth of 33,334 reads/cell. The CD34^−^GYPA^+^ (CD34N-CD235AP) sample generated 5,957 cells at a depth of 42,728 reads/cell. The CD34^+^ bulk (CD34P-Bulk) sample generated 10,082 cells at a depth of 29,308 reads/cell. The GPI-80^+^ (CD34N-GPI80P) sample generated 8,529 cells at a depth of 33,017 reads/cell. Quality control was performed with the singleCellTK package (Jenkins, 2018). We observed 9904, 4901, 9870.5, 10187 median UMIs, and 2681, 1401, 2808, 2923 median genes detected for the samples CD34^−^GYPA^−^, CD34^−^GYPA^+^, CD34^+^ bulk, and GPI-80^+^, respectively (**Figure S6**). The median levels of contamination estimated by DecontX was less than 1% in all samples (Yang et al., 2020). For the combined analysis, the four samples were merged and then normalized using SCTransform. In this step cell degradation and cell-cycle effects were regressed out. To cluster cells principal component analysis (PCA) was performed and the top 30 principal components (PCs) were used as input for the Louvain clustering algorithm (resolution 0.75). For 2D visualization purposes the dimension reduction algorithm UMAP was ran on the top 30 PCs using default settings. Differential expression tests were done with MAST (Finak et al., 2015). The same analysis pipeline was used to analyze the recently published fetal liver cell atlas (Popescu et al., 2019). In addition, the top 20 markers for each of the cell types described in Popescu et al. (2019) were computed using a Wilcoxon test and ranked by Log-fold-change. The corresponding gene signatures for each cell type were used to score their enrichment in our four samples, enabling the annotation of our dataset based on the modules extracted from the Fetal Liver Atlas. The correspondence between cell types present in both datasets was also validated by integrating both datasets using the pipeline previously mentioned and a harmonization step to correct batch effects. A cluster representing cells with high mitochondrial content and mixed lineage identities was excluded from downstream analysis. Six clusters identified in the progenitor compartment were grouped into one HSC/MPP cluster for the representation in Figure 1A. For the separate analysis of the CD34+ HSC/MPP compartment, clusters with a majority of CD34^+^ cells were extracted from the merged analysis of all four samples and re-clustered in an analogous manner as described above with resolution = 0.5. From this subset, cells were separated by original identity into a GPI80^+^ fraction and a CD34^+^ bulk fraction for downstream analyses, maintaining the cluster identities established upon reclustering of the CD34+ HSC/MPP compartment.

##### Trajectory analysis

Following the transcriptomic analysis described above of the four combined samples, we performed trajectory analysis in R version 3.6 (Stuart et al., 2019) using a trajectory analysis package: Monocle version 3.2 (Cao et al., 2019; Levine et al., 2015; McInnes, 2018; Qiu et al., 2017; Traag et al., 2019; Trapnell et al., 2014). We used the UMAP embeddings/coordinates from the transcriptomic analysis to perform trajectory analysis. Branches of the trajectory were created using clustering information and cell lineage determinations from the transcriptomic analysis. Each branch consists of the HSC/MPP populations and the clusters from one of the four lineages (erythroid cells, lymphoid B cells, T and NK cells, and myeloid cells), which were analyzed separately as four separate trajectories. A root node was selected for each branch from within the HSC/hematopoietic progenitor cell populations using the unbiased method provided in the Monocle 3 documentation (https://cole-trapnell-lab.github.io/monocle3/). The cells in each branch were assigned a pseudotime value with the Monocle 3 pseudotime ordering algorithms and the root node set as time zero. Gene expression changes over pseudotime were investigated in each branch and selected lineage markers were highlighted using heatmaps.

##### Comparison of LMNA expression

In order to investigate *LMNA* expression in prenatal HSPCs versus postnatal HSPCs we obtained raw count data from publicly available data sets (Belluschi et al., 2018; Velten et al., 2017; Zheng et al., 2018). These three data sets were merged with the single sample analysis of the CD34^+^ bulk sample and then processed using the same steps described for the transcriptomic analysis of our four samples combined in order to compare expression of *LMNA.*

#### CITE-seq: ADT processing and analysis

##### ADT data transformation and background removal

Non-specific staining using oligo-tagged antibodies was minimal and was corrected based on background staining levels detected in mouse cells (mESCs) that were spiked into samples to account for noise. ADT centered-log-ratio (CLR) transformation and background removal was performed as reported with adaptations (Stoeckius et al., 2017). Specifically, for 8735 CD34^+^bulk cells and 207 mouse cell spike-ins, raw ADT count data were CLR-transformed for each of the ADTs, using the function NormalizeData with normalization.method = “CLR” and margin = 1 in Seurat V3.2.1. The value at one standard deviation greater than the average CLR-transformed ADT counts from mouse cells were defined as the background cutoff and were subtracted from the 8735 human CD34^+^bulk cells. The same procedure was applied to 7235 GPI-80^+^ cells and 188 mouse cell spike-ins. The background-removed ADT data were used for all downstream analysis including UMAP, dot plots coloring and in silico gating.

##### Dimension reduction using UMAP

Log-normalized mRNA data and CLR-transformed ADT data were grouped into 10 bins and colored according to their feature expression levels respectively. For CLR-transformed ADT data, the top 1 percent feature values were labeled outlier and removed by specifying max.cutoff = “q99” in FeaturePlot function in Seurat V3.

#### In silico gating strategy

For *in silico* sorting purposes, cells were first gated based on CD34^+^CD38^−^ criteria and subsequently underwent signature-specific gating (schematic overview presented in **Figure S5A**). This process was guided by the average percentages obtained for the different marker combinations in flow cytometry experiments on 5 individual FL samples (**Figure S5A-B**). Because the CD38 oligo-tagged antibody in our CITE-seq panel did not result in ADT signal due to technical issues specific to this antibody, we based CD38^−^ selection in the first gating on mRNA values for CD38. While in this study we used a GPI-80 selection step to enrich for engraftment potential resulting in the profiling of 7235 GPI-80^+^ cells and establishment of a detailed engraftable FL HSC signature, an oligo-tagged GPI-80 antibody was not included in our CITE-seq panel. Given the poor correlation between GPI-80 protein expression and the corresponding mRNA expression of its encoding gene *VNN2* (**Figure S5C**), we consulted the extended transcriptomic signature of the GPI-80^+^ cell population to derive a GPI-80 module score and approximate GPI-80 protein expression in the CD34^+^CD38^−^ gated population. This module score was obtained by generating an AddModuleScore based on the top 30 enriched genes within the GPI-80^+^ HSC/MPP fraction and used to retrospectively label putative GPI-80 expressing cells within the CD34^+^CD38^−^ population based on how strongly their expression profile resembled GPI-80^+^ sorted cells (Tirosh et al., 2016). The module scores were calculated using the AddModuleScore function in Seurat V3.2.1 with features being the top 30 positively enriched genes within the GPI-80^+^ HSC/hematopoietic progenitor cell fraction, assay = “SCT”, and default parameters. When displaying the resulting continuous ‘GPI-80 module score’ values versus CD34+ ADT values, we found a distribution that strongly resembled FACS data (**Figure S5D**). To approximate what was sorted for the CITE-seq analysis based on GPI-80 cell surface marker expression, the top 2.37% of CD34^+^ cells positive for GPI-80 identity based on this GPI-80 score were considered (corresponding to 3% of CD34^+^CD38^−^ gated cells, **Figure S5A**). Those GPI-80 score^+^ cells that were also CD133^+^, were gated to reflect the Sumide et al. signature for comparison to the ‘classical’ (CD90^+^CD49f^+^) and ‘EPCR^+^’ (CD201^+^) signature, which were both gated based on ADT expression values.

For two-dimensional density plots showing in silico gating thresholds. Density was estimated using two-dimensional kernel density estimation with an axis-aligned bivariate normal kernel, evaluated on a square grid. Function kde2d in R package MASS V7.3-51.6 was used with number of grid points in each direction equals 500 (n = 500).

#### Processing of PBMCs

Mononuclear cells were isolated from a Leukoreduction System (LRS) Chamber (New York Biologics Inc, M, age 65) over a Ficoll gradient (Cytiva, 17144003).

#### Multi-dimensional flow cytometry

All antibodies were purchased from commercial vendors (**Table S5**) and were pre-titrated using PBMC and FL cells. For the 22-color analysis, PBMC and FL cells were thawed, washed, blocked with Human FcBlock (BioLegend, 422301) and stained with Live Dead Blue amine dye (Thermo Fisher, L34961). A cocktail of antibodies was prepared fresh and supplemented with Monocyte Blocker (BioLegend, 426102) and Brilliant Buffer Plus (BD Biosciences, 566385). Cells were stained in the dark at 4°C and washed twice. A sub-panel of brightly-expressed markers was assessed as a separate stain to serve as a fluorescence-minus-multiple control for further data interpretation. PBMC cells and single stain Ultracomp beads (Thermo Fisher, 01-2222-42) were stained to provide single stain spectral controls. Cells and single stain controls were analyzed on a 5-laser Aurora spectral flow cytometry (Cytek Biosciences) and raw fluorescence data from 64 channels were unmixed using ordinary least square algorithm in Spectraflo v2 (Cytek Biosciences).

#### Fluorescence cytometry data analysis

Data analysis pipelines were built using cloud-based OMIQ analysis platform (Omiq). Briefly, single cell data were asinh transformed (cofactor 6000) and 100,000 (as defined by least size FL sample) live single cell events from each sample were selected for analysis. All fluorescence data (excluding live-dead dye staining intensity) from 5 FL samples and 1 PBMC sample were projected into two-dimensional space with UMAP algorithm (McInnes, 2018) (neighbors = 15; minimum distance = 0.4; learning rate = 1; epochs = 200). For analysis, intensities of each fluorescence parameter were overlaid on UMAP maps to represent expression levels. Same dataset was clustered with Phenograph (Levine et al., 2015) (k=20, distance metric = euclidean) and clusters were color-coded and overlaid over UMAP maps. Phenograph clusters were also plotted as heatmaps to represent cell abundance over multiple clusters and median fluorescence intensity across multiple surface protein markers.

### QUANTIFICATION AND STATISTICAL ANALYSIS

Specifics about the replicates used in each experiment are available in the figure legends or specified in the text. The p value threshold to determine significance was set at p = 0.05. p value annotations on graphs are as follows: *p < 0.05, **p < 0.01 and are based on unpaired, two-tailed Student’s t tests. Data for quantitative experiments is typically represented as the mean with error bars representing the standard error of the mean, as specified in the figure legends.

### ADDITIONAL RESOURCES

The CITE-seq dataset described in this work has been made available in an interactive format at https://engraftable-hsc.cells.ucsc.edu

