## Supplemental Text and Figures for "Multi-Modal Profiling of Human Fetal Liver-Derived Hematopoietic Stem Cells Reveals the Molecular Signature of Engraftment Potential"

#### SUPPLEMENTAL FIGURE LEGENDS

**Figure S1. Experimental outline and gene expression changes over pseudotime**

(A) Overview of fractions isolated from the FL sample and how they were subdivided in individual samples to be run on the 10X Genomics platform for CITE-seq analysis and simultaneous transplantation experiments. Like the $CD34^+$ fraction, the $CD34^-$ flowthrough was first gated to select live cells prior to GYPA gating. The GPI-$80^+CD34^-$ cells captured in the GPI-80 sort gate represent macrophages. This population was not considered for the in-depth profiling of the GPI-$80^+$ fraction as this analysis starts from cells captured within the $CD34^+$ HSC/MPP fraction in **Figure 1A**.

(B-E) Pseudotime trajectories per hematopoietic lineage (left) and heatmaps illustrating gene expression changes over pseudotime for HSC/MPP and key lineage markers (right) for the erythroid (B), lymphoid (B cell) (C), lymphoid (T/NK cell) (D) and myeloid (E) lineage.

**Figure S2. Single cell transcriptomic profiling of the engraftment potential of FL-derived HSCs**

(A) Stacked barplot illustrating cell cycle distribution (percentage) per GPI-$80^+$ cluster. Cluster legend is shown in right upper corner of this Figure.

(B) Stacked barplot illustrating cell cycle distribution (percentage) per sample.

(C) Dot plot illustrating the distribution of a selection of genes enriched in the GPI-$80^+$ compared to $CD34^+$ bulk HSCs/MPPs. Expression is represented as z-scores. 'Percent expressed' represents the fraction of cells having non-zero expression values. Cluster legend is shown in right upper corner of this Figure.

(D) Comparison of *LMNA* expression in prenatal HSPCs versus postnatal HSPCs. We compared $CD34^+$ bulk FL HSC/MPPs from this study with existing scRNAseq data sets representing postnatal $CD34^+$ fractions (Zheng et al., 2018, Belluschi et al., 2018, Velten et al., 2017) in terms of *LMNA* expression. The expression range was divided in 'no expression', 'low expression' (bottom 75% *LMNA* expressing cells) and 'high expression' (top 25% *LMNA* expressing cells).

(E) EnrichR (https://maayanlab.cloud/Enrichr) read-out (-log(p-value)) for HSC/MPP cluster 2 in the GPI-80$^+$ sample. Black bars indicate when the FDR is significant with a score of <0.05.

(F) EnrichR (https://maayanlab.cloud/Enrichr) read-out (-log(p-value)) for HSC/MPP cluster 3 in the GPI-80$^+$ sample. Black bars indicate when the FDR is significant with a score of <0.05.

**Figure S3. Antibody-derived tags (ADTs) link cell surface protein expression to transcriptomic characterization of FL HSCs**

(A) Comparison of mRNA and ADT expression for a selection of HSC markers in the CD34$^+$bulk sample.

(B) Comparison of mRNA and ADT expression for a selection of HSC markers in the GPI-80$^+$ sample. The scale bar shows scaled expression from 0-Max.

(C-E) Transcriptomic cluster composition (%) within the CD34$^+$ bulk fraction upon stepwise enrichment for cell surface marker expression of CD90 (C), CD49f (D) and ENG (E). The x-axis is divided in segments showing the indicated top x% of expression for each marker.

**Figure S4. Multi-dimensional flow cytometric characterization across multiple FL samples**

(A) Phenograph analysis showing the distribution of clusters identified in **Figure 4D** over the different FL samples as well as PBMCs. Heat map color indicates percentage of cells per sample allocated to each cluster.

(B) Phenograph analysis representing clusters specific to the CD34$^+$ FL cells. The different cell surface markers assessed in this panel are shown on top. Heat map color indicates median fluorescence intensity of each marker expression per cluster and is normalized per column/marker.

(C) Overview of the contribution of each FL sample to the combined data set.

**Figure S5. In silico sorting strategy**

(A) Outline of *in silico* gating steps as guided by flow cytometry results. The left column shows the initial CD34$^{+}$CD38$^{-}$ gating step. The middle column shows the signature-specific gating including the percentages of cells that were gated based on ADT expression data (and GPI-80 score, see methods section). The right column shows representative FACS plots for a FL sample as well as the average percentage of cells in each of the gates based on flow cytometric analysis of 5 individual FL samples (<x%>, n=5).

(B) Box plots showing the distribution of cells in each gate based on flow cytometric analysis of 5 individual FL samples. The line within the box plot represents the average and the X represents the median.

(C) VNN2 mRNA expression in the GPI-80$^{+}$ sample

(D) FACS data (left) compared to a similar representation of the CITE-seq data (right) based on ADT expression for CD34 (y-axis) and the GPI-80$^{+}$ score (x-axis) (see methods section). One outlier cell with very high CD34 ADT expression (> 6) was excluded from the CITE-seq data (right).

**Figure S6. Data quality of CITE-seq experiment**

(A) Violin plots showing the total number of gene counts per cell for the samples CD34$^{-}$GYPA$^{-}$, CD34$^{-}$GYPA$^{+}$, CD34$^{+}$ bulk, and GPI-80$^{+}$. Red dotted lines indicate medians.

(B) Violin plots showing the total number of detected genes per cell for the four samples.

(C) Violin plots showing the levels of contamination estimated by DecontX for the four samples.

**Table S1. Oligo-tagged antibodies used in CITE-Seq**

The antibodies included in this panel consist of known markers relevant to characterize HSCs and other candidates identified in a pilot experiment.

**Table S2. DEGs between GPI-80$^{+}$ and CD34$^{+}$ bulk HSCs/MPPs**

**Table S3. DEGs between clusters within the GPI-80$^{+}$ sample**

**Table S4. Top enriched genes per cluster in the different HSC/ MPP fractions in Figures 2C-E.**

The rank is indicated behind the genes and the bolded gene names were selected for labeling of the different clusters in **Figure 2E**.

**Table S5. Multi-dimensional flow cytometry panel antibodies**

### Figure S1

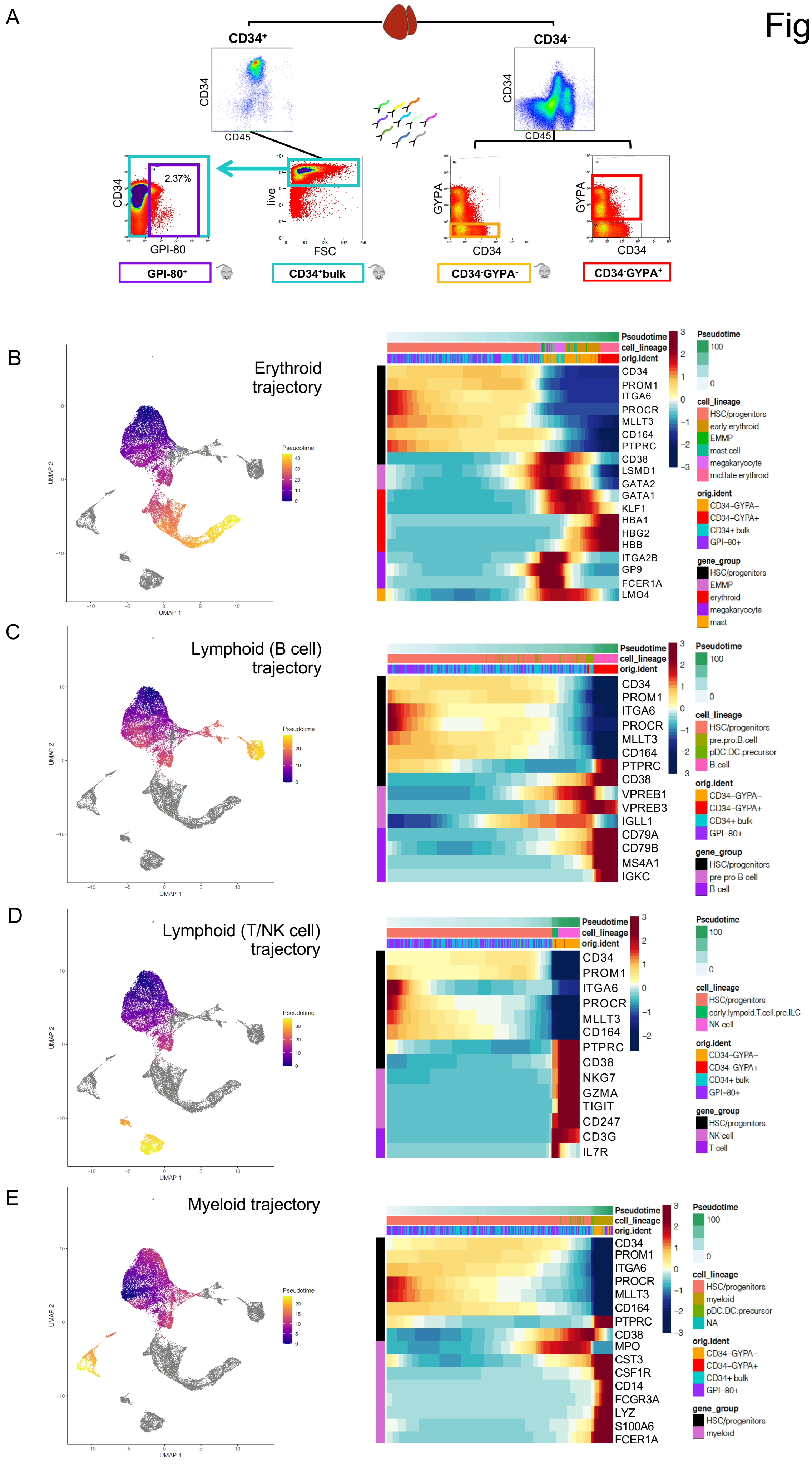

A
CD34+
CD34-
CD34
CD45
2.37%
GPI-80
live
FSC
GYPA
GPI-80+
CD34+bulk
CD34-GYPA-
CD34-GYPA+
B
Erythroid trajectory
C
Lymphoid (B cell) trajectory
D
Lymphoid (T/NK cell) trajectory
E
Myeloid trajectory
UMAP 1
UMAP 2
Pseudotime
cell_lineage
orig.ident
gene_group
HSC/progenitors
early.erythroid
EMMP
mast.cell
megakaryocyte
mid.late.erythroid
CD34−GYPA−
CD34−GYPA+
CD34+ bulk
GPI−80+
erythroid
mast
pre.pro.B.cell
pDC.DC.precursor
B.cell
pre pro B cell
B cell
early.lympoid.T.cell.pre.ILC
NK.cell
NK cell
T cell
myeloid
NA
CD34
PROM1
ITGA6
PROCR
MLLT3
CD164
PTPRC
CD38
LSMD1
GATA2
GATA1
KLF1
HBA1
HBG2
HBB
ITGA2B
GP9
FCER1A
LMO4
VPREB1
VPREB3
IGLL1
CD79A
CD79B
MS4A1
IGKC
NKG7
GZMA
TIGIT
CD247
CD3G
IL7R
MPO
CST3
CSF1R
CD14
FCGR3A
LYZ
S100A6

### Figure S2

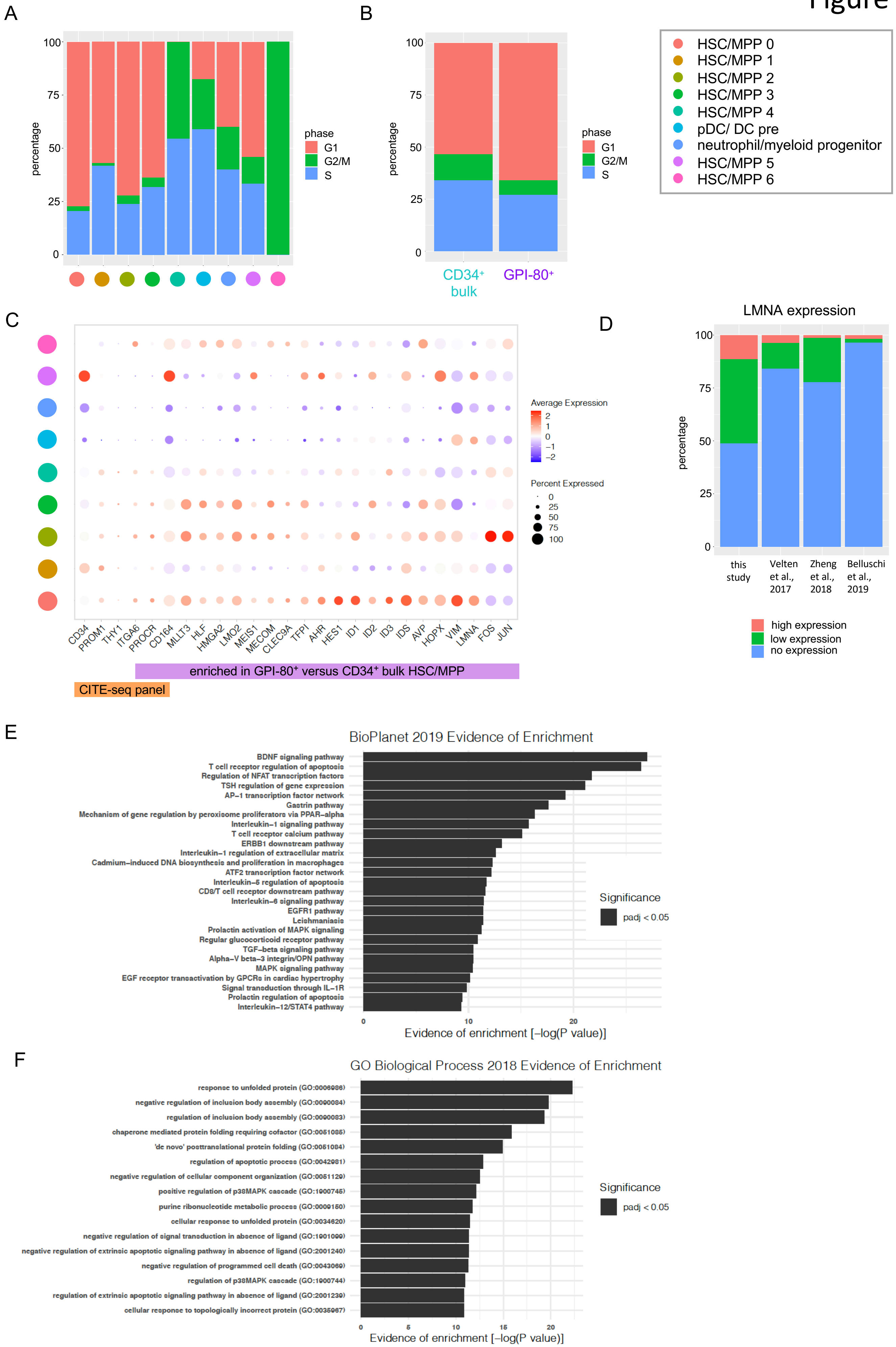
A
B
C
D
E
F
percentage
phase
G1
G2/M
S
CD34+ bulk
GPI-80+
HSC/MPP 0
HSC/MPP 1
HSC/MPP 2
HSC/MPP 3
HSC/MPP 4
pDC/ DC pre
neutrophil/myeloid progenitor
HSC/MPP 5
HSC/MPP 6
Average Expression
Percent Expressed
CD34
PROM1
THY1
ITGA6
PROCR
CD164
MLLT3
HLF
HMGA2
LMO2
MEIS1
MECOM
CLEC9A
TFPI
AHR
HES1
ID1
ID2
ID3
IDS
AVP
HOPX
VIM
LMNA
FOS
JUN
enriched in GPI-80+ versus CD34+ bulk HSC/MPP
CITE-seq panel
LMNA expression
this study
Velten et al., 2017
Zheng et al., 2018
Belluschi et al., 2019
high expression
low expression
no expression
BioPlanet 2019 Evidence of Enrichment
BDNF signaling pathway
T cell receptor regulation of apoptosis
Regulation of NFAT transcription factors
TSH regulation of gene expression
AP−1 transcription factor network
Gastrin pathway
Mechanism of gene regulation by peroxisome proliferators via PPAR−alpha
Interleukin−1 signaling pathway
T cell receptor calcium pathway
ERBB1 downstream pathway
Interleukin−1 regulation of extracellular matrix
Cadmium−induced DNA biosynthesis and proliferation in macrophages
ATF2 transcription factor network
Interleukin−5 regulation of apoptosis
CD8/T cell receptor downstream pathway
Interleukin−6 signaling pathway
EGFR1 pathway
Leishmaniasis
Prolactin activation of MAPK signaling
Regular glucocorticoid receptor pathway
TGF−beta signaling pathway
Alpha−V beta−3 integrin/OPN pathway
MAPK signaling pathway
EGF receptor transactivation by GPCRs in cardiac hypertrophy
Signal transduction through IL−1R
Prolactin regulation of apoptosis
Interleukin−12/STAT4 pathway
Significance
padj < 0.05
Evidence of enrichment [−log(P value)]
GO Biological Process 2018 Evidence of Enrichment
response to unfolded protein (GO:0006986)
negative regulation of inclusion body assembly (GO:0090084)
regulation of inclusion body assembly (GO:0090083)
chaperone mediated protein folding requiring cofactor (GO:0051085)
'de novo' posttranslational protein folding (GO:0051084)
regulation of apoptotic process (GO:0042981)
negative regulation of cellular component organization (GO:0051129)
positive regulation of p38MAPK cascade (GO:1900745)
purine ribonucleotide metabolic process (GO:0009150)
cellular response to unfolded protein (GO:0034620)
negative regulation of signal transduction in absence of ligand (GO:1901099)
negative regulation of extrinsic apoptotic signaling pathway in absence of ligand (GO:2001240)
negative regulation of programmed cell death (GO:0043069)
regulation of p38MAPK cascade (GO:1900744)
regulation of extrinsic apoptotic signaling pathway in absence of ligand (GO:2001239)
cellular response to topologically incorrect protein (GO:0035967)
Significance
padj < 0.05
Evidence of enrichment [−log(P value)]

Figure S3

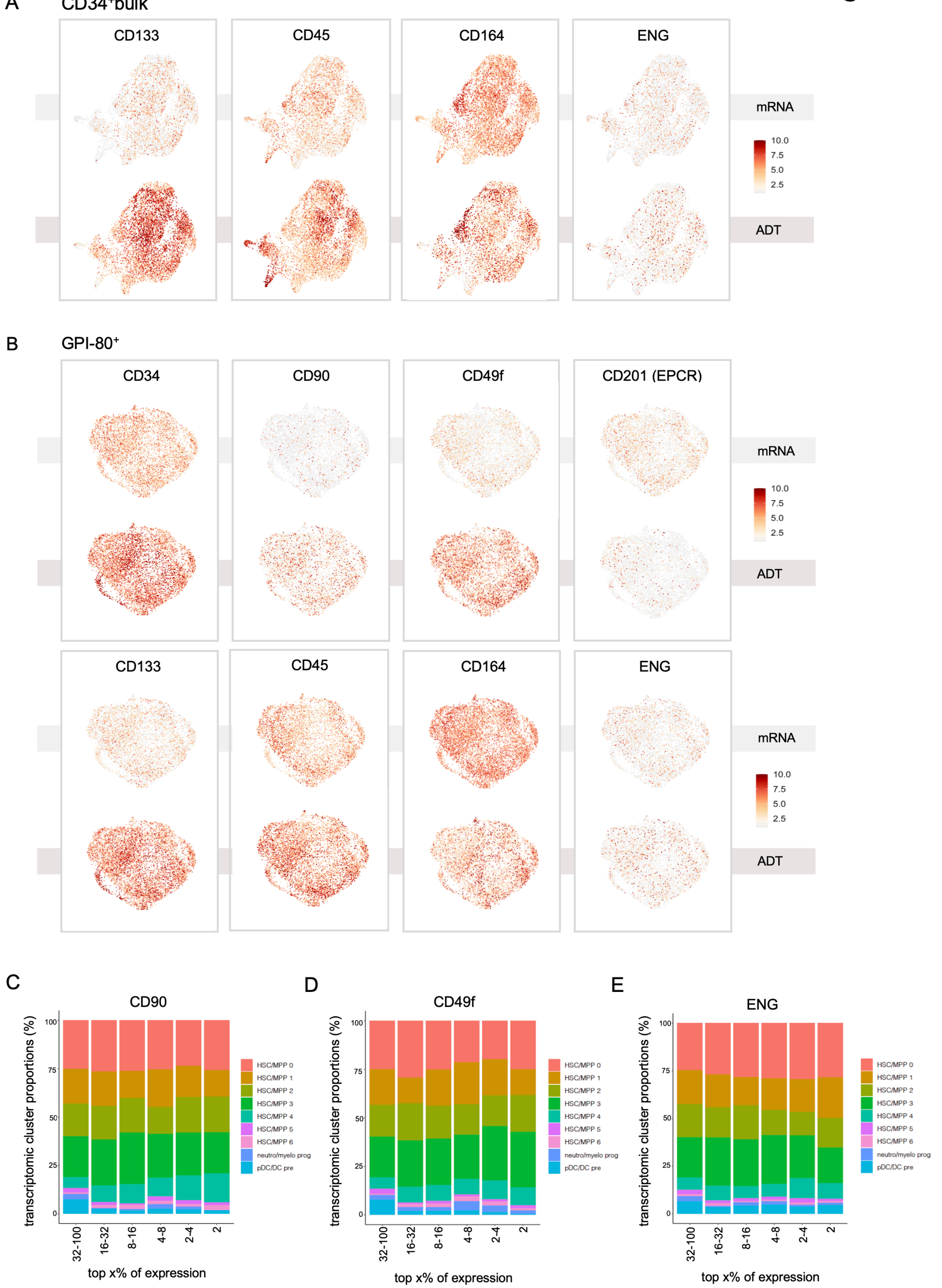
A
CD34⁺bulk
CD133
CD45
CD164
ENG
mRNA
ADT
10.0
7.5
5.0
2.5
B
GPI-80⁺
CD34
CD90
CD49f
CD201 (EPCR)
CD133
CD45
CD164
ENG
mRNA
ADT
C
CD90
D
CD49f
E
ENG
transcriptomic cluster proportions (%)
top x% of expression
32-100
16-32
8-16
4-8
2-4
2
0
25
50
75
100
HSC/MPP 0
HSC/MPP 1
HSC/MPP 2
HSC/MPP 3
HSC/MPP 4
HSC/MPP 5
HSC/MPP 6
neutro/myelo prog
pDC/DC pre

### Figure S4

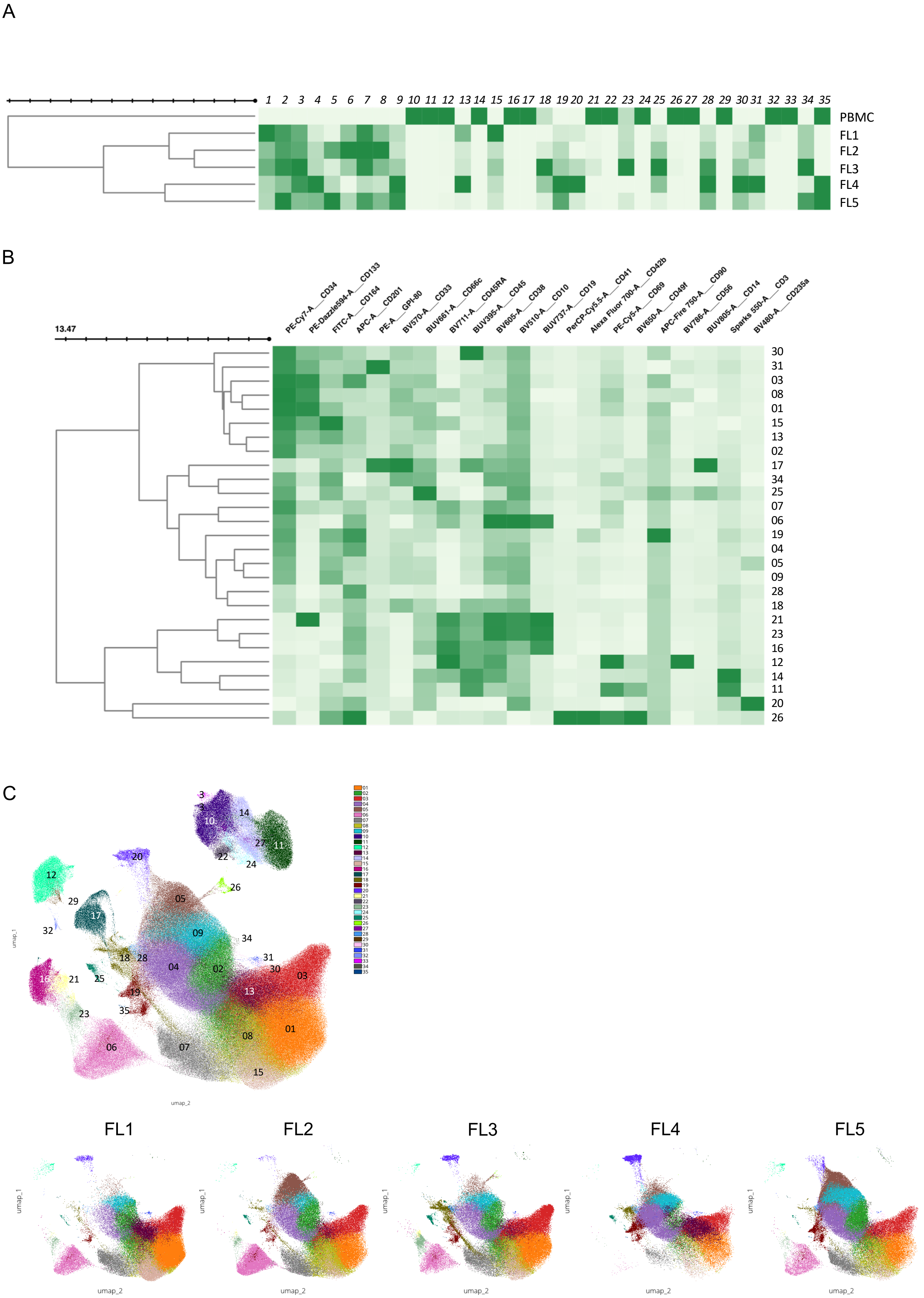
A
PBMC
FL1
FL2
FL3
FL4
FL5
B
13.47
PE-Cy7-A__CD34
PE-Dazzle594-A__CD133
FITC-A__CD164
APC-A__CD201
PE-A__GPI-80
BV570-A__CD33
BUV661-A__CD66c
BV711-A__CD45RA
BUV395-A__CD45
BV605-A__CD38
BV510-A__CD10
BUV737-A__CD19
PerCP-Cy5.5-A__CD41
Alexa Fluor 700-A__CD42b
PE-Cy5-A__CD69
BV650-A__CD49f
APC-Fire 750-A__CD90
BV786-A__CD56
BUV805-A__CD14
Sparks 550-A__CD3
BV480-A__CD235a
C
umap_1
umap_2
FL1
FL2
FL3
FL4
FL5

### Figure S5

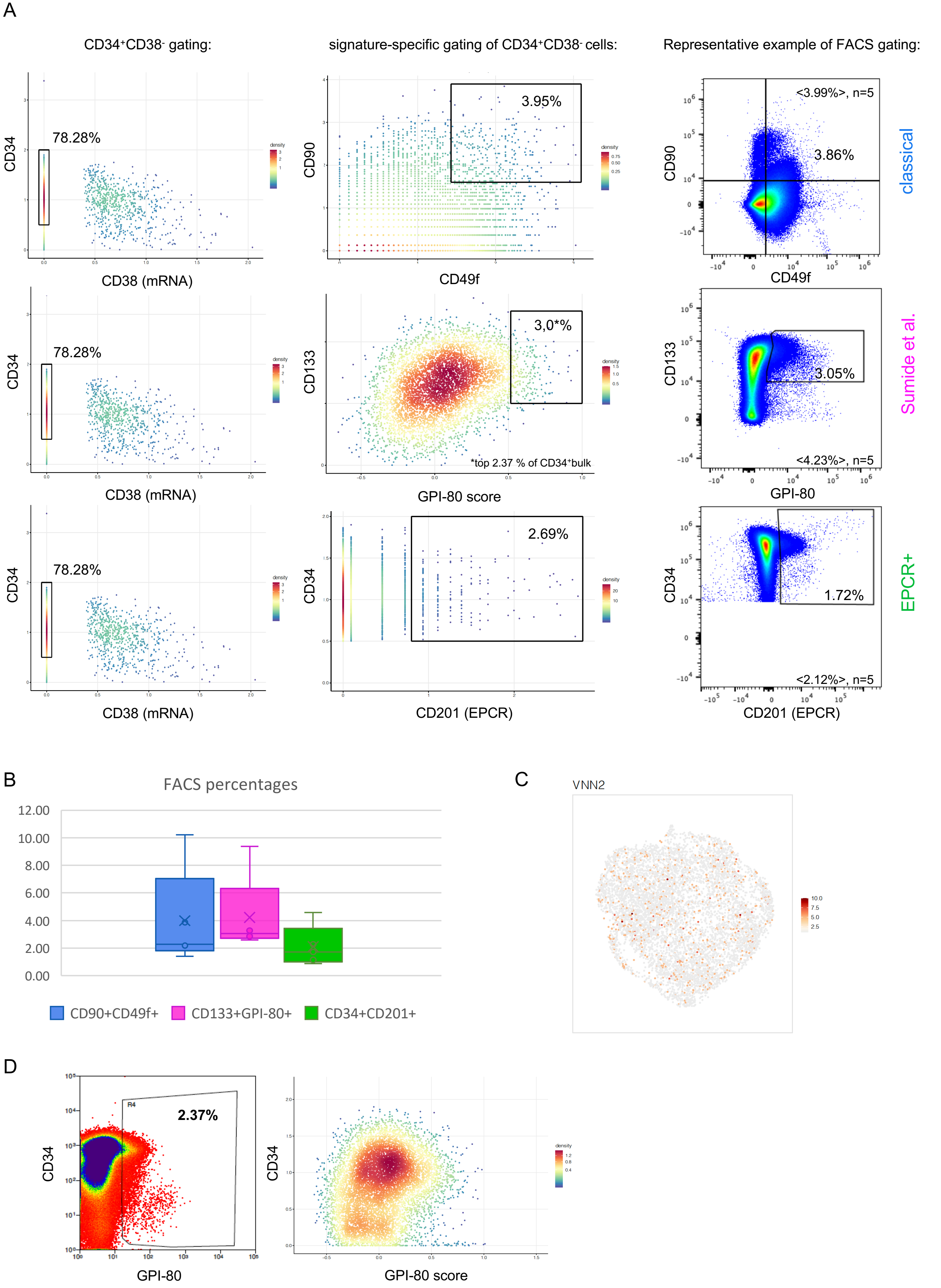
A
CD34+CD38- gating:
signature-specific gating of CD34+CD38- cells:
Representative example of FACS gating:
CD34
78.28%
CD38 (mRNA)
CD90
3.95%
CD49f
<3.99%>, n=5
3.86%
classical
CD133
3,0*%
*top 2.37 % of CD34+bulk
GPI-80 score
3.05%
<4.23%>, n=5
GPI-80
Sumide et al.
2.69%
CD201 (EPCR)
1.72%
<2.12%>, n=5
EPCR+
B
FACS percentages
12.00
10.00
8.00
6.00
4.00
2.00
0.00
CD90+CD49f+
CD133+GPI-80+
CD34+CD201+
C
VNN2
10.0
7.5
5.0
2.5
D
R4
2.37%
CD34
GPI-80
GPI-80 score

### Figure S6

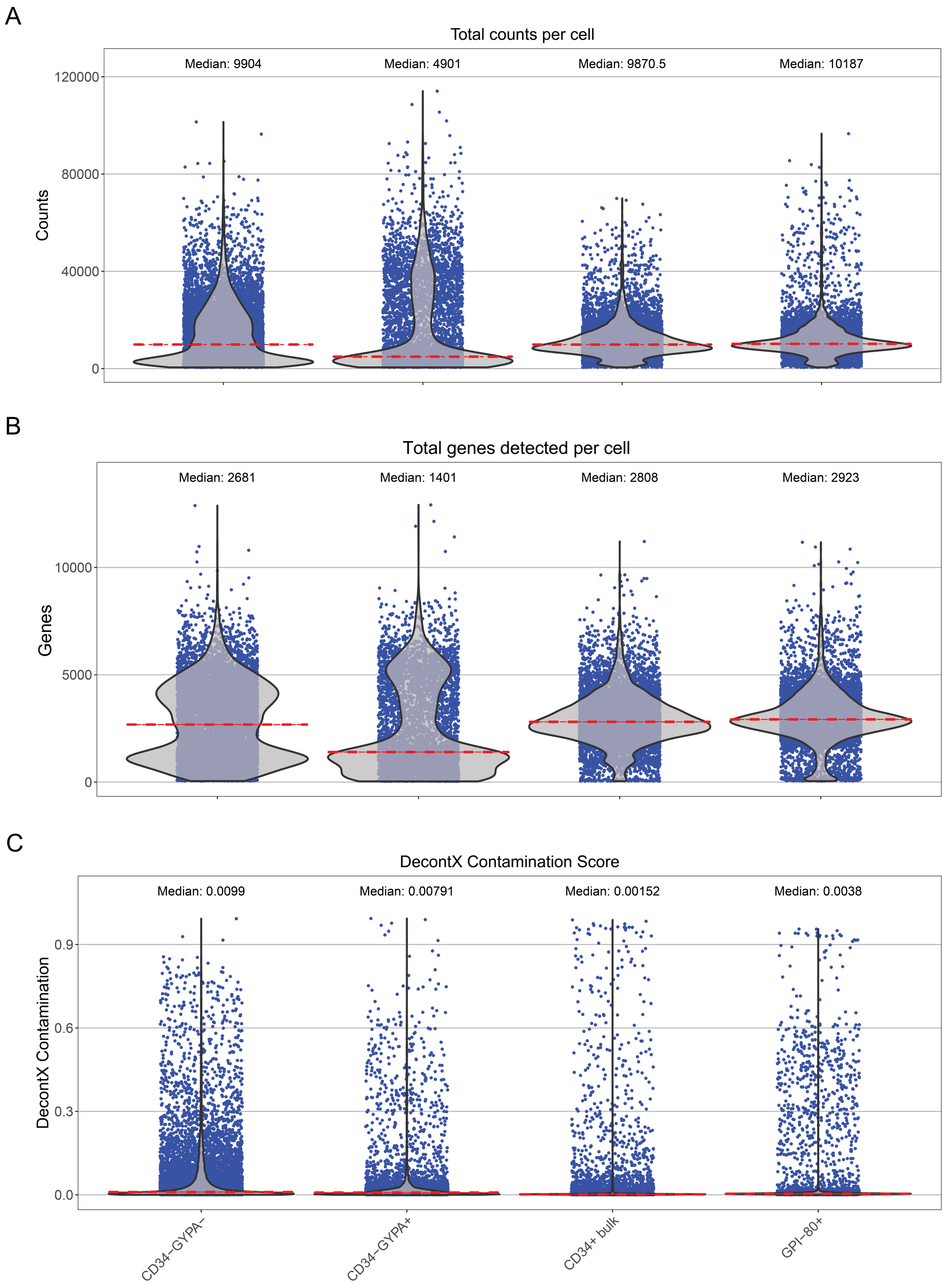
A
Total counts per cell
Median: 9904
Median: 4901
Median: 9870.5
Median: 10187
Counts
120000
80000
40000
0
B
Total genes detected per cell
Median: 2681
Median: 1401
Median: 2808
Median: 2923
Genes
10000
5000
0
C
DecontX Contamination Score
Median: 0.0099
Median: 0.00791
Median: 0.00152
Median: 0.0038
DecontX Contamination
0.9
0.6
0.3
0.0
CD34−GYPA−
CD34−GYPA+
CD34+ bulk
GPI−80+

**Table S1. Oligo-tagged antibodies used in CITE-Seq**

| Cell surface marker | Gene | Vendor | Catalog number |
|---|---|---|---|
| CD34 | *CD34* | BioLegend | 343537 |
| CD38 | *CD38* | BioLegend | 303541 |
| CD90 | *THY1* | BioLegend | 328135 |
| CD45 | *PTPRC* | BioLegend | 304064 |
| CD49f | *ITGA6* | BioLegend | 313633 |
| CD133 | *PROM1* | BioLegend | 372815 |
| CD201 | *EPCR/PROCR* | BioLegend | 351907 |
| CD164 | *CD164* | BioLegend | 324809 |
| CD105 | *ENG* | BioLegend | 323221 |
| CD144 | *CDH5/ VECAD* | BioLegend | 348517 |
| CD202b | *TEK/TIE2* | BioLegend | 334213 |
| VEGFR3 | *FLT4* | BioLegend | 356207 |
| CLEC1B | *CLEC1B* | BioLegend | 372009 |
| CD31 | *PECAM1* | BioLegend | 303137 |
| CD184 | *CXCR4* | BioLegend | 334213 |
| HLA-DR,DP,DQ | *HLA class II genes* | BioLegend | 307659 |
| pan-HLA class I | *HLA class I genes* | BioLegend | 311445 |
| CD141 | *THBD* | BioLegend | 344121 |
| CD235a | *GYPA* | BioLegend | 334213 |

**Table S4. Top enriched genes in HSC/MPP clusters across samples**

| cluster | HSC/MPP (Figure 2C) | CD34+ bulk (Figure 2D) | GPI-80+ (Figure 2E) |
|---|---|---|---|
| HSC/MPP 0 | ID1 (1), VIM (2), RGCC (3), **LMNA (4)** | RGCC (1), **LMNA (2)** | RGCC (1), **LMNA (2)** |
| HSC/MPP 1 | **CDK6 (1)** | IGLL1 (1), HCST (2), **CDK6 (3)** | IGLL1 (1), HCST (2), **CDK6 (3)** |
| HSC/MPP 2 | **FOS (1), JUN (2)** | **FOS (1), JUN (2)** | **FOS (1), JUN (2)** |
| HSC/MPP 3 | HSPA1A (1), **HSPA1B (2)** | HSPA1A (1), **HSPA1B (2)** | **HSPA1B (1)**, HSPA1A (2) |
| HSC/MPP 4 | **HIST1H4C (1)** | **HIST1H4C (1)** | **HIST1H4C (1)** |
| pDC/ DC pre | **IGLL1 (1)**, LTB (2), IGJ (3) | **IGLL1 (1)**, LTB (2), IGJ (3) | **IGLL1 (1)**, LTB (2), IGJ (3) |
| neutrophil/myeloid progenitor | **MPO (1)**, PRTN3 (2), AZU1 (3), LFD (4), IGLL1 (5) | IGLL1 (1), **MPO (2)** | IGLL1 (1), **MPO (2)** |
| HSC/MPP 5 | **PRSS3P1 (1)** | **PRSS3P1 (1)** | **PRSS3P1 (1)** |
| HSC/MPP 6 | **CENPF (1)**, TOP2A (2) | **CENPF (1)**, TOP2A (2) | **CENPF (1)**, TOP2A (2) |

**Table S5. Multi-dimensional flow cytometry panel antibodies**

| marker | Fluorophore | Vendor | Catalog number |
|---|---|---|---|
| CD45 | BUV395 | BD Biosciences | 563791 |
| Live/dead | fixable UV live/dead | Thermo Fischer | L34961 |
| CD66c | BUV661 | BD Biosciences | 741653 |
| CD19 | BUV737 | BD Biosciences | 612757 |
| CD14 | BUV805 | BD Biosciences | 612902 |
| CD235a | Pacific Blue | BD Biosciences | 746358 |
| CD10 | BV510 | BioLegend | 312219 |
| CD33 | BV570 | BioLegend | 303417 |
| CD38 | BV605 | BioLegend | 303532 |
| CD49f | BV650 | BD Biosciences | 563706 |
| CD45RA | BV711 | BioLegend | 304137 |
| CD56 | BV786 | BD Biosciences | 564058 |
| CD164 | FITC | BioLegend | 324805 |
| CD3 | Spark Blue 550 | BioLegend | 100259 |
| GPI-80 | PE | MBL International | D087-5 |
| CD133 | PE-Dazzle 594 | BioLegend | 372811 |
| CD41 | PerCP-Cy55 | BioLegend | 303719 |
| CD69 | PE-Cy5 | BioLegend | 310907 |
| CD34 | PE-Cy7 | BioLegend | 343515 |
| CD201 (EPCR) | APC | BioLegend | 351906 |
| CD42b | Alexa 700 | BioLegend | 303927 |
| CD90 | APC-Fire750 | BioLegend | 328137 |
